# VAMP7-dependent mitochondria-lysosome contacts contribute to glial mitochondrial dynamics and dopaminergic neuron survival

**DOI:** 10.1101/2021.08.16.456582

**Authors:** Honglei Wang, Mengxiao Wang, Yu-Ting Tsai, Yi-Hsuan Cheng, Yu-Tung Lin, Chia-Ching Lin, Yi-Hua Lee, Linfang Wang, Shuanglong Yi, Shiping Zhang, Yufeng Pan, Margaret S. Ho

## Abstract

Disrupted mitochondrial dynamics in neurons are linked to neurodegenerative diseases, but their regulation in glia remains poorly understood. Here, we show that the R-SNARE protein VAMP7 regulates the untethering of mitochondria–lysosome contacts (MLCs) in adult fly glia. Glial-specific knockdown of VAMP7 leads to prolonged MLCs and mitochondrial elongation related to altered fission/fusion dynamics. These VAMP7-deficient mitochondria exhibit hyperpolarized membrane potential, resulting in increased ROS activity, lipid droplet accumulation, and dopaminergic (DA) neuron degeneration. Mechanistically, VAMP7 interacts with the GTPase-activating protein TBC1D15-17 to promote Rab7 GTP hydrolysis. Without VAMP7, TBC1D15-17 remains bound to Rab7 but fails to activate its hydrolysis, resulting in elevated GTP-bound Rab7 and impaired MLCs untethering. Consistently, expression of GTP-locked Rab7^Q67L^ or GAP-dead TBC1D15-17^ΔGAP^ phenocopies the mitochondrial defects, while GDP-bound Rab7^T22N or wild-type TBC1D15-17 restores the MLC dynamics. Considering that SNARE proteins mediate membrane fusion, our results demonstrate a new role for VAMP7 in glial mitochondrial dynamics via organelle contacts, impacting neuron survival in a non-cell-autonomous manner.

## Introduction

Mitochondria are dynamic organelles that undergo structural and functional changes in response to a variety of cellular signals such as energy production, metabolism, or survival. They constantly remodel and interact with other organelles so that their morphologies are swiftly modified to adapt the cellular needs. Different aspects of mitochondrial dynamics have been uncovered including fission (division), fusion, or shape transition, generating different forms of mitochondria that function in various cellular contexts^1, 2, 3^. These dynamic changes in morphology and function are under tight regulation and have been linked to catastrophic consequences such as neurodegeneration and death^4, 5^. For instance, mitochondrial fission regulated by the dynamin-related protein-1 (Drp1), a GTPase that dynamically associates with the endoplasmic reticulum (ER) and mitochondria^6, 7, 8, 9^, not only occurs in healthy mitochondria to accommodate physiological needs, but also suppresses oxidative damage and safeguards neuron survival^10^. It is generally considered that the fission process generates smaller mitochondria easier to be targeted by mitophagy, the autophagosomal degradation of damaged mitochondria mediated by the Parkinson’s disease risk genes *PINK1* and *Parkin*^11, 12, 13^. On the other hand, elongated mitochondria lacking fission accumulate oxidative damage and lose respiratory function, transforming into large spheres resistant to mitophagy. Interestingly, most of these mitochondrial dynamic defects underlying neurodegeneration are ascribed to neurons, whereas how mitochondrial dynamics in the brain cells “glia” contribute to neurodegeneration remains poorly understood.

In addition to ER, mitochondrial fission sites are also contacted by lysosomes, which help to restrict the mitochondrial motility by recruiting the lysosomal GTPase Rab7^14, 15^. It has been shown that Rab7 GTP hydrolysis, converting the Rab7-GTP to the inactive GDP form, is crucial for the untethering of lysosome and mitochondria, marking the site for mitochondrial fission^15^. Rab7 GTP hydrolysis is regulated by guanine nucleotide exchange factors (GEFs) that stimulate the exchange of bound GDP for GTP, resulting in Rab7 activation, and GTPase activating proteins (GAPs) that facilitate Rab7 GTP hydrolysis, thereby inactivating Rab7^16^. These findings implicate a role for lysosome in regulating mitochondrial dynamics. In regard to this aspect, whether lysosome and its related factors play a role in glial mitochondrial fission is largely unknown; how mitochondrial dynamic is controlled in glia has not been extensively studied.

The *Drosophila* vesicular R-SNARE (soluble N-ethylmaleimide-sensitive factor attachment protein receptors), VAMP7, is homologous to mammalian VAMP7 and VAMP8, and the latter exhibits a prominent expression in microglia^17, 18^. Mammalian VAMP7 or VAMP8 localizes to the late endosome and lysosome and forms a complex with the cytosolic Synaptosomal-associated protein 29 (SNAP-29) and the autophagosomal Syntaxin-17 (STX17). Together, this complex regulates the autophagosome-lysosome fusion in autophagy^19, 20^. Despite a role in vesicle fusion, whether VAMP7 plays additional roles with respect to its lysosomal origin remains unclear. Considering the lysosomal impact on mitochondrial dynamics and a possible role of VAMP7 in glia, it is worthy of investigating whether the lysosomal VAMP7 regulates mitochondrial dynamics in glia. Our findings reveal that VAMP7 regulates the untethering of mitochondria-lysosome contacts (MLCs) in adult fly glia. Reduced VAMP7 expression in glia causes fission/fusion-dependent mitochondrial elongation, likely owing to the persistent and prolonged MLCs. These elongated mitochondria are dysfunctional with hyperpolarized membrane potential, increased ROS levels, and the production of lipid droplets, ultimately leading to dopaminergic (DA) neurodegeneration in the brain. Mechanistically, VAMP7 interacts with the Rab7 GAP TBC1D15-17, facilitating Rab7 GTP hydrolysis to control the dynamics of MLCs. Considering that SNARE proteins mediate membrane fusion, our findings uncover a new role for a canonical SNARE factor in glial mitochondrial dynamics, demonstrating its importance in glial contribution to neuron survival.

## Results

### Adult fly glia deficient of VAMP7 exhibit mitochondrial elongation

To investigate VAMP7 function in glia, two independent fly RNAi lines: *vamp7-*RNAi^38300^ and *vamp7-*RNAi^43543^ were used to silence *vamp7* expression in adult fly glia. To assess the efficacy of these lines, 3-day-old adult fly brains were first collected to analyze *vamp7* mRNA levels upon RNAi expression in glia. RT-qPCR analyses revealed a significant decrease in *vamp7* mRNA levels, with *vamp7-*RNAi^43543^ causing a greater reduction (Figure S1A). To examine *vamp7* mRNA levels only in adult fly glia, 100 adult fly brains expressing *UAS-mCD8.GFP*, with or without *vamp7-*RNAi, under the control of the pan-glial driver *repo*-GAL4 were dissected and sorted by fluorescence-activated cell sorting (FACS). Consistently, RT-qPCR analyses revealed significantly decreased *vamp7* mRNA levels in sorted glia upon expressing *vamp7-*RNAi^43543^ (Figure S1B). Hence, *vamp7-*RNAi^43543^ was used as the major RNAi line throughout the study.

We then investigated the consequences of silencing glial *vamp7* expression in the brains by Transmission Electron Microscopy (TEM). A marker *UAS-HRP* was expressed in glia (*repo>HRP*), attempting to label the glial processes visualized in black. Mitochondria were identified for their typical structures of inner and outer mitochondrial membranes, with the inner membranes folding as cristae and extending into the matrix (Figure 1A). Compared to the 3-day-old control brains expressing the *Luciferase* RNAi (*luc*-RNAi), mitochondria were elongated when expressing *vamp7-*RNAi^43543^ in glia, indicating a possible defect of fission or fusion (enclosed by white dashed lines, Figure 1A). The value of mitochondrial length over diameter (L/D) was significantly increased (Figure 1B). In addition, the ratio of mitochondria with a L/D number lower than 2.5 (L/D<2.5/Total) decreased, while the ratio of those with a L/D number higher than 2.5 (L/D>2.5/Total) increased upon glial VAMP7 reduction (Figure 1C).

**Figure 1.**
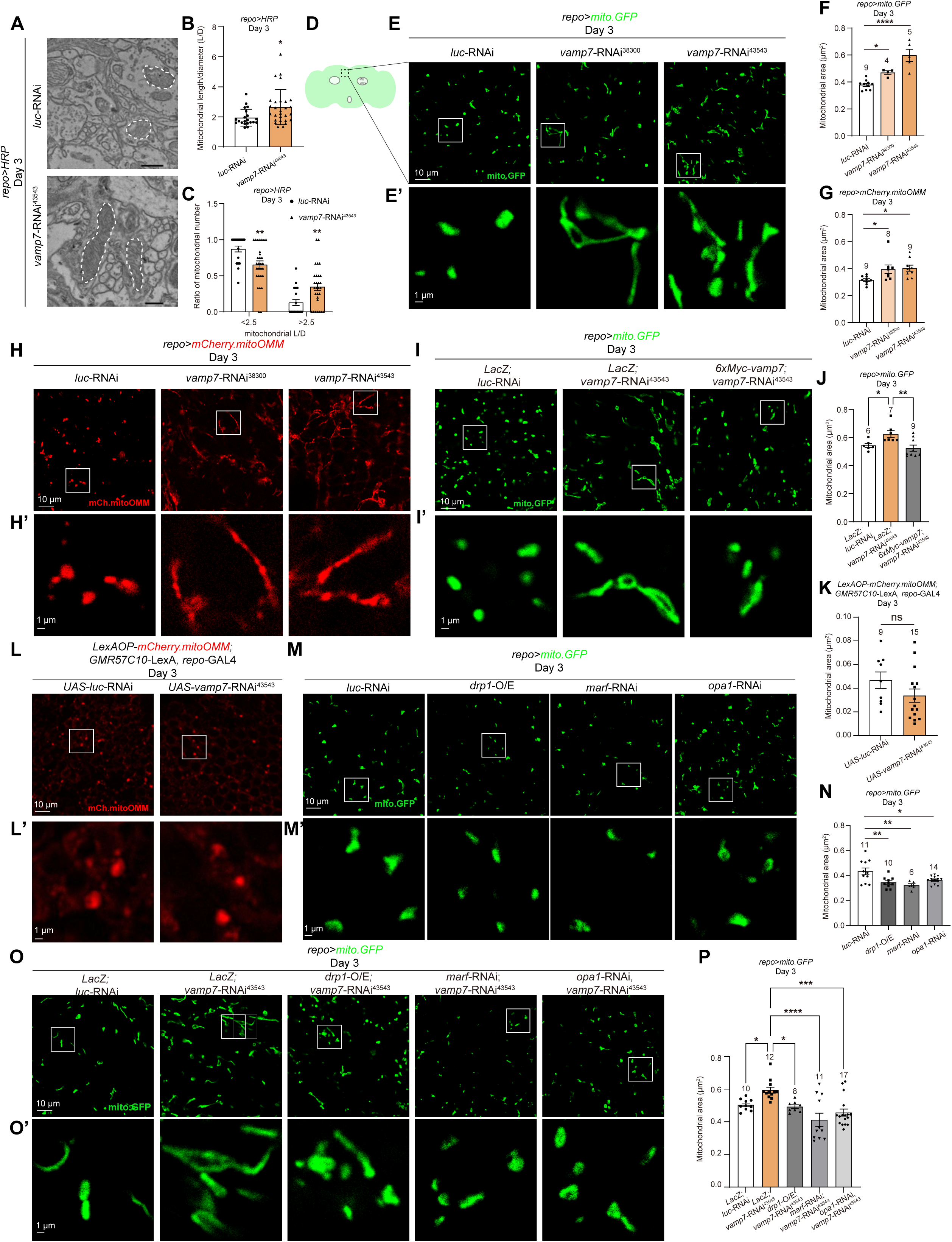
Mitochondrial elongation in VAMP7-deficient adult fly glia. (**A-C**) Representative TEM images (A) of mitochondria (double-membraned with cristae) in control and *repo*>*vamp7*-RNAi^43543^ adult fly brains. *UAS-HRP* was also expressed in glia, attempting to label the glial processes in black. Note the presence of elongated mitochondria (white dashed lines) upon glial VAMP7 reduction. Quantifications (B and C) of mitochondrial morphology are designated as mitochondrial length/diameter (L/D, B) and the ratio of mitochondria with a L/D number higher or lower than 2.5 over the total number (C). Note that the ratio of mitochondria with L/D number greater than 2.5 is significantly higher upon glial VAMP7 reduction. (**D**) Illustration of an adult fly brain with symmetrical mushroom body calyces located posteriorly (green). Dashed square in black indicates the dorsal-medial region where images throughout the study were acquired. (**E**-**H**) Representative images (E and H) and quantifications (F and G) of mitochondrial morphology labeled by *UAS-mito.GFP* (E and E’) or *UAS-mCherry.mitoOMM* (H and H’) in control and *vamp7*-RNAi^43543^-expressing adult fly glia. Note that the average mitochondrial area quantified by ImageJ in 2D is significantly larger than the control glia when expressing *vamp7*-RNAi^43543^. (**I**-**L**) Representative images (I and L) and quantifications (J and K) of mitochondrial morphology labeled by *UAS-mito.GFP* (I and I’) or *LexAOP-mCherry.mitoOMM* (L and L’) in control and *vamp7*-RNAi^43543^-expressing adult fly glia. Note that Co-expression of 6xMyc-*vamp7* suppresses the elongation (I and J). Neuronal mitochondria were analyzed in adult fly brains expressing *LexAOP-mCherry.mitoOMM* under the pan-neuronal driver *GMR57C10*-LexA and *UAS-vamp7*-RNAi^43543^ by *repo*-GAL4. Note that neuronal mitochondria remain largely intact upon glial VAMP7 reduction. (**M-P**) Representative images (M,M’,O,O’) and quantifications (N and P) of glial mitochondria labeled by *UAS-mito.GFP* (*repo>mito.GFP*) upon expression of the fission/fusion factors Drp1 (*drp1*-O/E), *marf*-RNAi, or *opa1*-RNAi with or without VAMP7. Note that increasing expression of the fission factor Drp1 or downregulating the expression of the fusion factor Marf or Opa1 causes increased fission, opposite to the mitochondrial elongation phenotype when silencing *vamp7* expression in glia. Co-expression of *drp1*-O/E, *marf*-RNAi, or *opa1*-RNAi with *vamp7*-RNAi^43543^ in glia suppresses mitochondrial elongation. The selected white squares in E,H,I,L,M,O are enlarged and aligned in E’,H’,I’,L’,M’,O’. All samples examined are brains of 3-day-old adult flies unless noted otherwise. For single transgene experiment, *Luciferase*-RNAi (*luc*-RNAi) was used as a control. For double transgene experiment, *UAS-LacZ*; *luc*-RNAi was used as a control. Scale bars of different sizes are indicated on the images (TEM=500 nm). For TEM, all mitochondria found in more than 20 brain regions were analyzed per sample. Mitochondrial length, diameter, and area are quantified by ImageJ as detailed in the Materials and Methods. Serial confocal Z-stack sections were taken at similar planes across all genotypes, showing representative single-layer images. Statistical graphs are shown with scatter dots and the number of brains or mitochondria listed on top. For TEM, one scatter dot represents the average of mitochondria L/D or ratio analyzed. Data are shown as mean ± SEM. P-values of significance (indicated with asterisks, ns no significance, * p<0.05, ** p<0.01, *** p<0.001 and **** p<0.0001) are calculated by two-tailed unpaired t-test between two sample groups or by one-way ANOVA among three sample groups followed by Tukey’s multiple comparisons test.

To verify these results, two independent mitochondrial reporter lines: *UAS-mito.HA.GFP* (*mito.GFP*) and *UAS-mCherry.mitoOMM* (*mCherry.mitoOMM*), which express a mitochondrial import signal and label the outer mitochondrial membrane, respectively, were used to assess the mitochondrial morphology in glia silencing *vamp7* expression. Consistent with the EM results, glial VAMP7-depleted mitochondria were elongated with significantly increased area in 3-day-old adult fly brains (Figures 1D-1H). A consistent increase was also detected when using a different quantification method to assess the mitochondrial volume (Figures S1C and S1D, please see Methods for quantification details). For consistency, the dorsal-medial region of the adult fly brain was analyzed throughout the study. Interestingly, reintroducing VAMP7 by expressing a Myc-tagged *vamp7* transgene in glia suppressed the mitochondrial elongation upon silencing glial *vamp7* (Figures 1I and 1J). Interestingly, neuronal mitochondria labeled by expressing *LexAOP-mCherry.mitoOMM* under the control of the pan-neuronal driver *GMR57C10*-GAL4 remain largely intact with normal area and morphology when simultaneously expressing *vamp7-*RNAi^43543^ in glia (Figures 1K and 1L). Thus, these results indicate that VAMP7 regulates mitochondrial morphology in adult fly glia.

### Mitochondrial elongation in VAMP7-deficient adult fly glia depends on the expression of mitochondrial fission and fusion factors

Given that mitochondrial elongation is linked to altered fission or fusion, we next asked if glial VAMP7 genetically interacts with the fission factor Dynamin-related protein 1 (Drp1) or the fusion factors Mitochondrial assembly regulatory factor (Marf) and Optic atrophy 1 (Opa1). Consistent with previous findings in other cell types, Drp1 overexpression (*drp*-O/E, increased fission) or downregulated Marf or Opa1 expression by RNAi (*marf*-RNAi or *opa1*-RNAi, decreased fusion) in glia caused a reduction in the mitochondrial area, indicative of their function in fission and fusion (Figures 1M and 1N)^21, 22^. Interestingly, *drp*-O/E, *marf*-RNAi, or *opa1*-RNAi expression significantly suppressed the mitochondrial elongation in the *vamp7-*RNAi^43543^-expressing glia (Figures 1O and 1P), suggesting that altered fission/fusion helps prevent mitochondrial elongation induced by silencing *vamp7* expression in glia. These results suggest that VAMP7-mediated mitochondrial dynamics in adult fly glia are fission/fusion dependent.

### VAMP7 regulates the dynamics of mitochondria-lysosome contacts in adult fly glia

It has been reported that lysosomes contact mitochondria to restrict their motility and determine the fission site^15, 23^. Given the lysosomal localization of VAMP7 and the VAMP7-mediated mitochondrial elongation, we next investigated whether VAMP7 regulates mitochondrial morphology via the mitochondria-lysosome contacts (MLCs) in adult fly glia. Two reporters: *mCherry.mitoOMM* and *UAS-Lamp1.GFP* (*Lamp1.GFP*) were co-expressed to label mitochondria and lysosomes, respectively, so to visualize glial MLCs in live brains. Images acquired during the time-lapse analysis revealed that mCherry and GFP fluorescent puncta were in close contact and then dissociated in the control glia at t=120s (white arrow, Figure 2A and Movie S1). Whereas dissociation between the mCherry and GFP fluorescent puncta was also detected at t=100s in the control glia in a parallel experiment (white arrow, top panel in Figure 2B), persistent and prolonged contacts were observed in glia expressing either *vamp7-*RNAi, up to t=200s (middle and bottom panels in Figure 2B and Movies S2-S4). The frequency and duration of mCherry-positive mitochondria and GFP-positive lysosomes contacting each other increased significantly when reducing glial *vamp7* expression (Figures 2B-2D). Consistent with the observations on mitochondrial morphology, reintroducing VAMP7 in glia suppressed these contact deficits. Whereas MLCs in the control glia expressing two *UAS* transgenes (*LacZ*; *luc*-RNAi) began to dissociate at t=80s, MLCs in the *vamp7-*RNAi^43543^-expressing glia remained persistently associated (middle panel in Figure 2E). Upon VAMP7 co-expression in glia, MLCs dissociation was restored and occurred faster at t=170s (white arrows, bottom two panels in Figure 2E). The frequency and duration of these MLCs were also suppressed by VAMP7 co-expression (Figures 2E-2G and Movies S5-S7). Taken together, these results suggest that VAMP7 regulates the dynamics of MLCs in adult fly glia.

**Figure 2.**
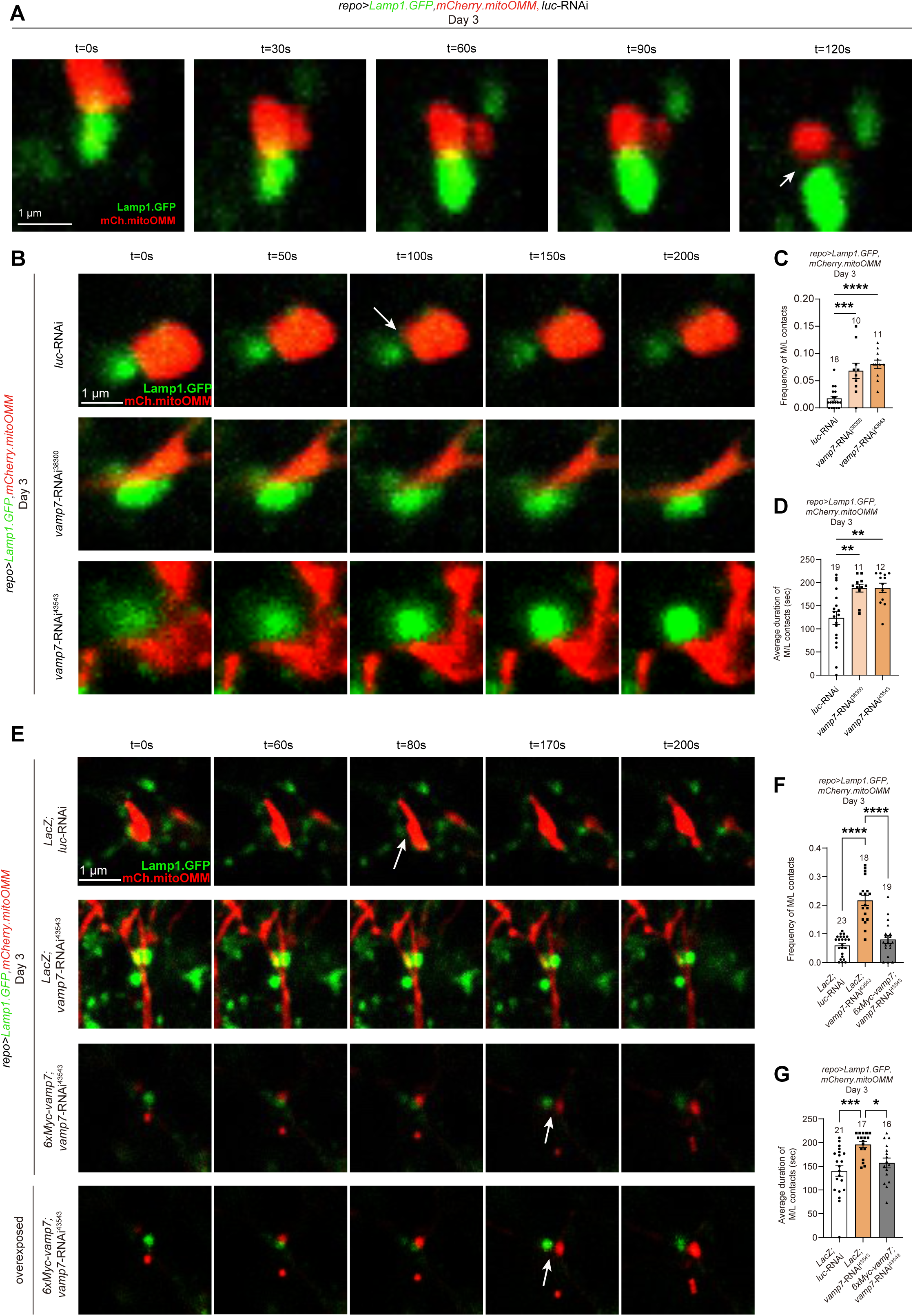
VAMP7 regulates the dynamics of mitochondria-lysosome contacts in adult fly glia. (**A-G**) Representative images (A,B,E) and quantifications (C,D,F,G) on *in-vivo* time lapse analysis of mitochondria-lysosome contacts (MLCs) in adult fly glia expressing two reporters: *UAS-mCherry.mitoOMM* (labels the mitochondrial outer membrane, red) and *UAS-Lamp1.GFP* (labels lysosome, green). In control glia, the contacts start untethered at t=120s (white arrow, A) or t=100s (white arrow in top panel, B), whereas the untethering of contacts is delayed up to t=200s in *vamp7*-RNAi^43543^-expressing glia (middle and bottom panels, B). Note that the frequency and duration of the contacts increase (C and D) in glia expressing either *vamp7*-RNAi. In a parallel experiment, 6xMyc-*vamp7* expression restores the contact dynamics (untethering at t=170s, white arrows in bottom two panels, E). An overexposed panel is shown for the rescue 6xMyc-*vamp7* genotype for clarity (bottom panel, E). Note that the frequency and duration of the contacts (F and G) are restored upon 6xMyc-*vamp7* co-expression in glia. All samples examined are brains of 3-day-old adult flies unless noted otherwise. For single transgene experiment, *luc*-RNAi was used as a control. For double transgene experiment, *UAS-LacZ*; *luc*-RNAi was used as a control. White arrows indicate the untethering. Scale bars of different sizes are indicated on the images. Serial confocal Z-stack sections were taken at similar planes across all genotypes, showing representative merged images. Statistical graphs are shown with scatter dots and the number of brains (one MLC per brain) listed on top. Images for the time lapse series are shown in 3x3 μm squares centering on the target MLC selected from the movies (Movies S1-S7). Parameters for the contact dynamics in C,D,F,G are quantified using Imaris and ImageJ. Data are shown as mean ± SEM. P-values of significance (indicated with asterisks, ns no significance, * p<0.05, ** p<0.01, *** p<0.001 and **** p<0.0001) are calculated by one-way ANOVA followed by Tukey’s multiple comparisons test.

### GTP-bound Rab7 levels increase upon silencing glial *vamp7* expression

Previous studies have demonstrated that the untethering of MLCs depends on Rab7 GTP hydrolysis^15^. We next checked if VAMP7 regulates the dynamics of MLCs via Rab7 GTP hydrolysis. In light of the reported GST-Glutathione Transferase-Rab Interacting Lysosomal Protein (RILP) assay for measuring the GTP-bound Rab7 levels^24^, the gene encoding RILP in flies, *RILP-like* (*RILPL*), was fused with a N-terminal three tandem repeat HA tag (*3xHA-RILPL*) and cloned for transgenic expression in flies. RILPL preferentially binds to and specifically reflects GTP-bound Rab7 levels. By co-immunoprecipitation (Co-IP) analysis, 3-day-old adult fly brains expressing 3xHA-RILPL in glia were dissected and homogenized for further pull-downs using beads conjugated to the anti-HA antibodies. Validating the approach, expression of the constitutively active form of Rab7, 3xFlag-Rab7^Q67L^, resulted in increased levels of GTP-bound Rab7 as quantified by the ratio of Rab7 in the 3xHA-RILPL pull downs over total Rab7 in the input (Figures 3A and 3B). Notably, GTP-bound Rab7/total Rab7 ratio also increased upon glial VAMP7 reduction, suggesting that silencing VAMP7 disrupts Rab7 GTP hydrolysis, leading to increased GTP-bound Rab7 levels (Figures 3A and 3B).

**Figure 3.**
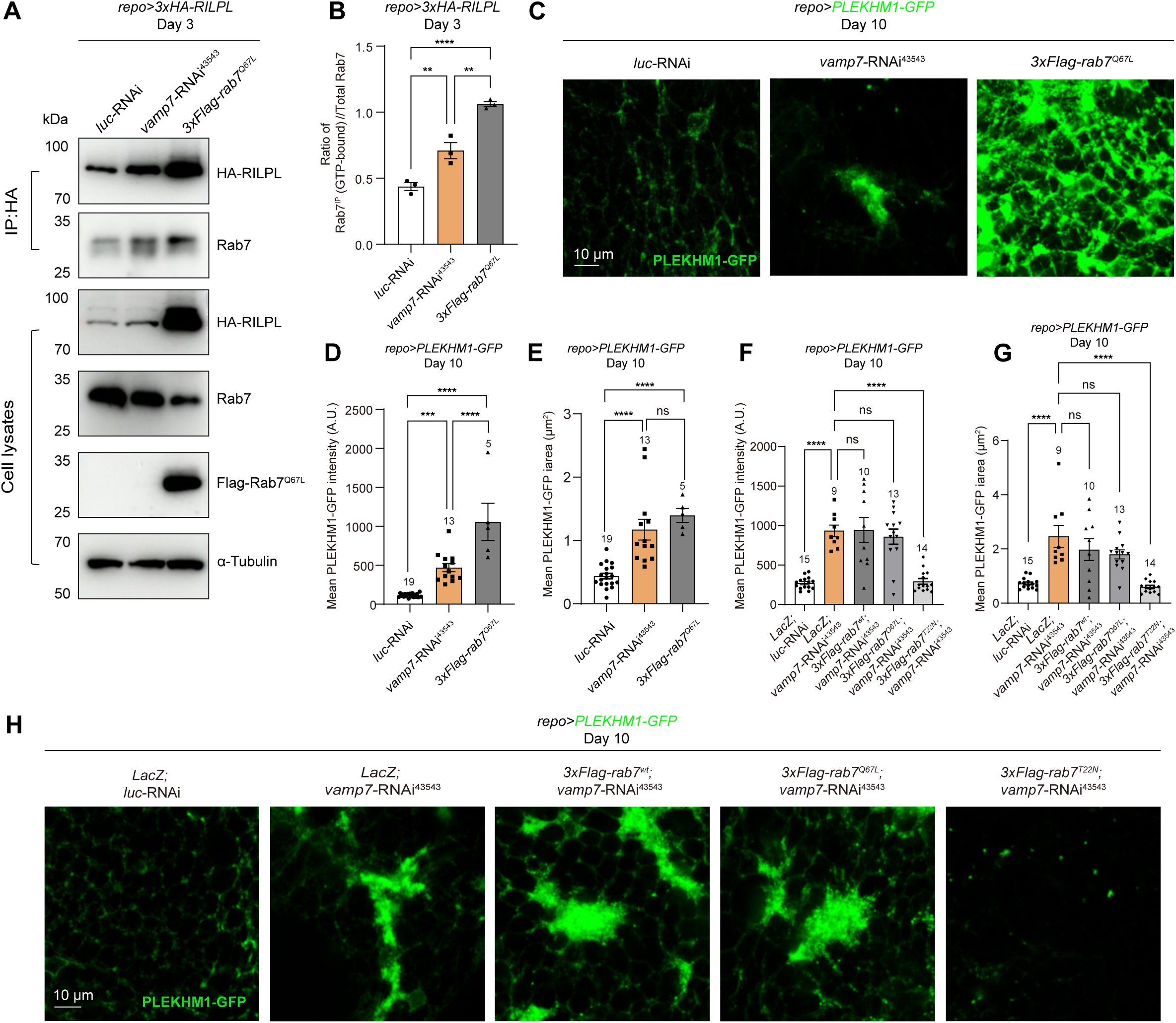
GTP-bound Rab7 levels increase upon glial VAMP7 reduction. (**A** and **B**) Representative WB images (A) and quantifications (B) of Rab7-RILPL interaction. GTP-bound Rab7 are present in the eluates of glial RILPL pull-downs by the HA-conjugated beads (*repo>3xHA-RILPL*, A). Note that GTP-bound Rab7 levels increases when silencing *vamp7* expression or overexpressing Rab7^Q67L^ in glia. (**C**-**H**) Representative images (C and H) and quantifications (D-G) of transgenic flies expressing PLEKHM1(CG6613)-GFP. Note that GFP signals exhibit aggregated pattern in forms of multiple puncta and elevated intensities upon glial VAMP7 reduction. All samples examined are brains of 3-day-old adult flies unless noted otherwise. For single transgene experiment, *luc*-RNAi was used as a control. For double transgene experiment, *UAS-LacZ*; *luc*-RNAi was used as a control. Scale bars of different sizes are indicated on the images. Serial confocal Z-stack sections were taken at similar planes across all genotypes, showing representative merged images. Statistical graphs are shown with scatter dots and the number of brains listed on top. For Co-IP analysis, 200 independent fly brains for each genotype were analyzed. Data are shown as mean ± SEM. P-values of significance (indicated with asterisks, ns no significance, * p<0.05, ** p<0.01, *** p<0.001 and **** p<0.0001) are calculated by ordinary one-way ANOVA followed by Tukey’s multiple comparisons test.

Previously reported Rab7 effectors include RILP (used in the present study) and two other related proteins PLEKHM1 and Rubicon^25^. Of these, *Drosophila* PLEKHM1 (CG6613) was shown to interact with Rab7 in a GTP-dependent manner^26^. Based on these findings, transgenic flies expressing PLEKHM1-GFP were made for testing the GTP-bound Rab7 levels in flies. Consistent with the RILPL results, PLEKHM1-GFP levels increased as quantified by GFP intensities and area when expressing *vamp7-*RNAi^43543^ or 3xFlag-Rab7^Q67L^, indicating more GTP-bound Rab7 (Figures 3C-3E). Interestingly, co-expression of 3xFlag-Rab7^WT^ or the constitutively active 3xFlag-Rab7^Q67L^ fails to impact the increased GTP-bound Rab7 levels upon glial VAMP7 reduction, whereas co-expression of the GDP-bound 3xFlag-Rab7^T22N^ suppressed the increase (Figures 3F-3H). These results suggest that glial VAMP7 regulates Rab7 GTP hydrolysis as reflected by the level of two Rab7 effectors 3xHA-RILPL and PLEKHM1-GFP; supplying more GDP-bound Rab7 helps restore the Rab7 GTP/GDP balance *in-vivo*.

### VAMP7 regulates the dynamics of MLCs via Rab7 GTP hydrolysis in adult fly glia

Given that the untethering of MLCs depends on Rab7 GTP hydrolysis and glial VAMP7 regulates both the dynamics of MLCs and the GTP-bound Rab7 levels, we next asked if VAMP7 regulates the untethering of MLCs via Rab7 GTP hydrolysis. By conducting the *in-vivo* time lapse analysis using *mCherry.mitoOMM* and *Lamp1.GFP*, the glial MLCs were monitored under various genetic conditions (Figures 4A-4C and Movies S8-S12). The impaired untethering of MLCs was consistently detected in adult fly glia with reduced *vamp7* expression. Interestingly, only the co-expression of Rab7^T22N^, but not Rab7^WT^ nor Rab7^Q67L^, significantly restored the prolonged contact dynamics in the absence of glial VAMP7. As the control MLCs untethered at around t=180s (white arrow in top panel, Figure 4A), the frequency and duration of the contacts were restored when co-expressing Rab7^T22N^ and *vamp7-*RNAi^43543^ (t=140s, white arrows in bottom two panels, Figures 4A-4C). Taken together, these results suggest that increasing GDP-bound Rab7 levels restores the disrupted dynamics of MLCs upon glial VAMP7 reduction; VAMP7 regulates the balance of Rab7 GTP/GDP levels attributing to the untethering of the MLCs in adult fly glia.

**Figure 4.**
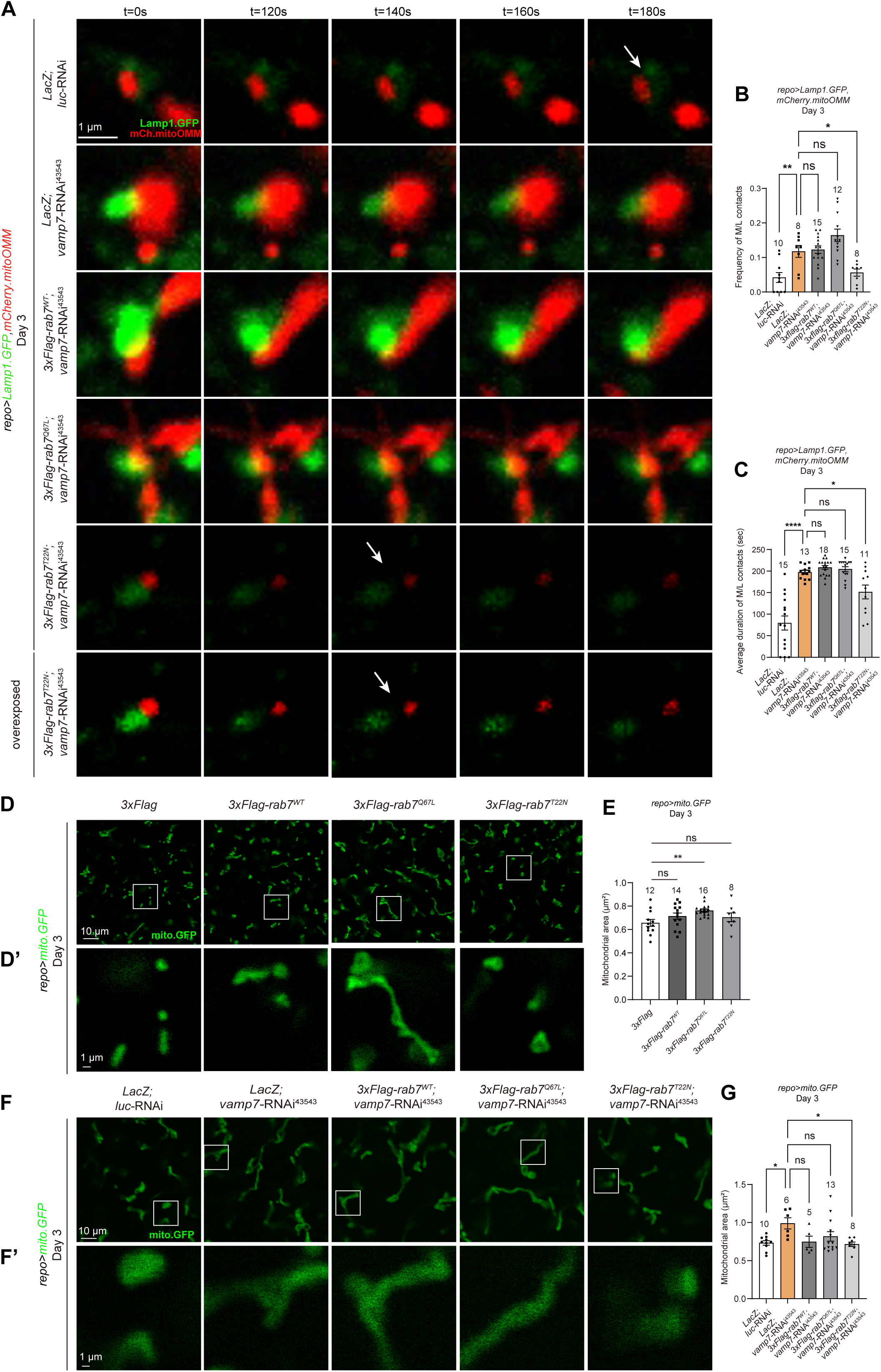
VAMP7 regulates mitochondria-lysosome contacts via Rab7 hydrolysis in adult fly glia. (**A-C**) Representative images (A) and quantifications (B and C) on *in-vivo* time lapse analysis of mitochondria-lysosome contacts (MLCs) in adult fly glia expressing two reporters: *UAS-mCherry.mitoOMM* (labels the mitochondrial outer membrane, red) and *UAS-Lamp1.GFP* (labels lysosome, green). Note that VAMP7-deficient contacts exhibit a higher frequency and prolonged duration (>180s), whereas the control contacts start untethering at t=180s (white arrow in top panel, A). Note that only the glial expression of the GDP-bound Rab7 (*3xFlag-Rab7^T22N^*), but not Rab7 (*3xFlag-Rab7^wt^*) nor GTP-bound Rab7 (*3xFlag-Rab7^Q67L^*), restores the contact dynamics (white arrows in bottom two panels in A, B and C). An overexposed panel is shown for the rescue *3xFlag-Rab7^T22N^* genotype for clarity (bottom panel, A). (**D**-**G**) Representative images (D,D’,F,F’) and quantifications (E and G) of mitochondrial morphology labeled by *UAS-mito.GFP* (*repo*>*mito.GFP*) in control, *3xFlag-Rab7^wt^-*, *3xFlag-Rab7^Q67L^-, 3xFlag-Rab7^T22N^-, or vamp7*-RNAi^43543^-expressing glia. Note that the average mitochondrial area quantified by ImageJ is significantly larger upon Rab7^Q67L^ expression in glia. The mitochondrial area is also restored when co-expressing Rab7^T22N^ and *vamp7*-RNAi^43543^. The selected white squares in D and F are enlarged and aligned in D’ and F’. All samples examined are brains of 3-day-old adult flies unless noted otherwise. For single transgene experiment, *luc*-RNAi was used as a control. For double transgene experiment, *UAS-LacZ*; *luc*-RNAi was used as a control. White arrows indicate the untethering. Scale bars of different sizes are indicated on the images. Serial confocal Z-stack sections were taken at similar planes across all genotypes, showing representative single layer or merged (MLCs) images. Statistical graphs are shown with scatter dots and the number of brains (one MLC per brain) listed on top. Images for the time lapse series are shown in 3x3 μm squares centering on the target MLC selected from the movies (Movies S8-S12). Parameters for the contact dynamics in B-D are quantified using Imaris and ImageJ. Data are shown as mean ± SEM. P-values of significance (indicated with asterisks, ns no significance, * p<0.05, ** p<0.01, *** p<0.001 and **** p<0.0001) are calculated by one-way ANOVA followed by Tukey’s multiple comparisons test.

As the untethering of MLCs underlies mitochondrial fission, we also analyzed mitochondrial morphology when altering Rab7 GTP hydrolysis when silencing *vamp7* expression in glia. Interestingly, expression of Rab7^Q67L^ (i.e., increased GTP-bound Rab7 levels), but not Rab7^WT^ nor the GDP-bound Rab7^T22N^, caused mitochondrial elongation in adult fly glia (Figures 4D and 4E). Furthermore, co-expression of Rab7^T22N^, but not Rab7^WT^ nor Rab7^Q67L^, restored the *vamp7* RNAi^43543^-induced mitochondrial elongation in glia (Figures 4F and 4G). These results are consistent with the observations on the untethering of the MLCs, suggesting that the balance of Rab7 GTP/GDP levels is crucial for VAMP7-mediated mitochondrial dynamics in adult fly glia.

### VAMP7 interacts with the Rab7 GAP TBC1D15-17

Given that VAMP7 regulates GTP-bound Rab7 levels, we next asked if VAMP7 interacts with Rab7 or its associated GTPase-activating protein (GAP) TBC1D15-17, a member of the TBC (Tre2/Bub2/Cdc16)-domain-containing protein family that facilitates Rab7 GTP hydrolysis. TBC1D15-17 is a fly protein homologous to mammalian TBC1D15 and TBC1D17 with overall 30% identity and 63% and 61% similarity, respectively^27^. Mammalian TBC1D15 has been shown to localize to mitochondria via binding with the outer mitochondrial membrane protein Fis1, then drives Rab7 GTP hydrolysis to untether the MLCs^15^. To test interaction among VAMP7, TBC1D15-17, and Rab7, transgenic flies expressing these proteins with different tags in tandem repeats were generated. These flies carried *UAS-3xHA-TBC1D15-17*, *UAS-6xMyc-vamp7*, or *UAS-3xFlag-Rab7* to express the corresponding proteins. Different combinations of these proteins were then co-expressed in flies for Co-IP analyses. Using antibodies targeting different tags, a strong interaction between VAMP7 and TBC1D15-17 was detected, but not between VAMP7 and Rab7 (Figures 5A and 5B). While a strong interaction was also detected between TBC1D15-17 and Rab7, the interaction remains unaffected and strong upon *vamp7*-RNAi^43543^ expression in glia (Figures 5C and 5D). These results suggest that VAMP7 physically interacts with TBC1D15-17 and both are within the same protein complex. It is feasible that VAMP7-TBC1D15-17 interaction might be a prerequisite for TBC1D15-17 to mediate Rab7 GTP hydrolysis for the untethering of MLCs. Notably, TBC1D15-17 remains associated with Rab7 upon glial VAMP7 reduction, suggesting that VAMP7 is not required for their physical interaction; rather, VAMP7 interaction with TBC1D15-17 might promote its GAP activity.

**Figure 5.**
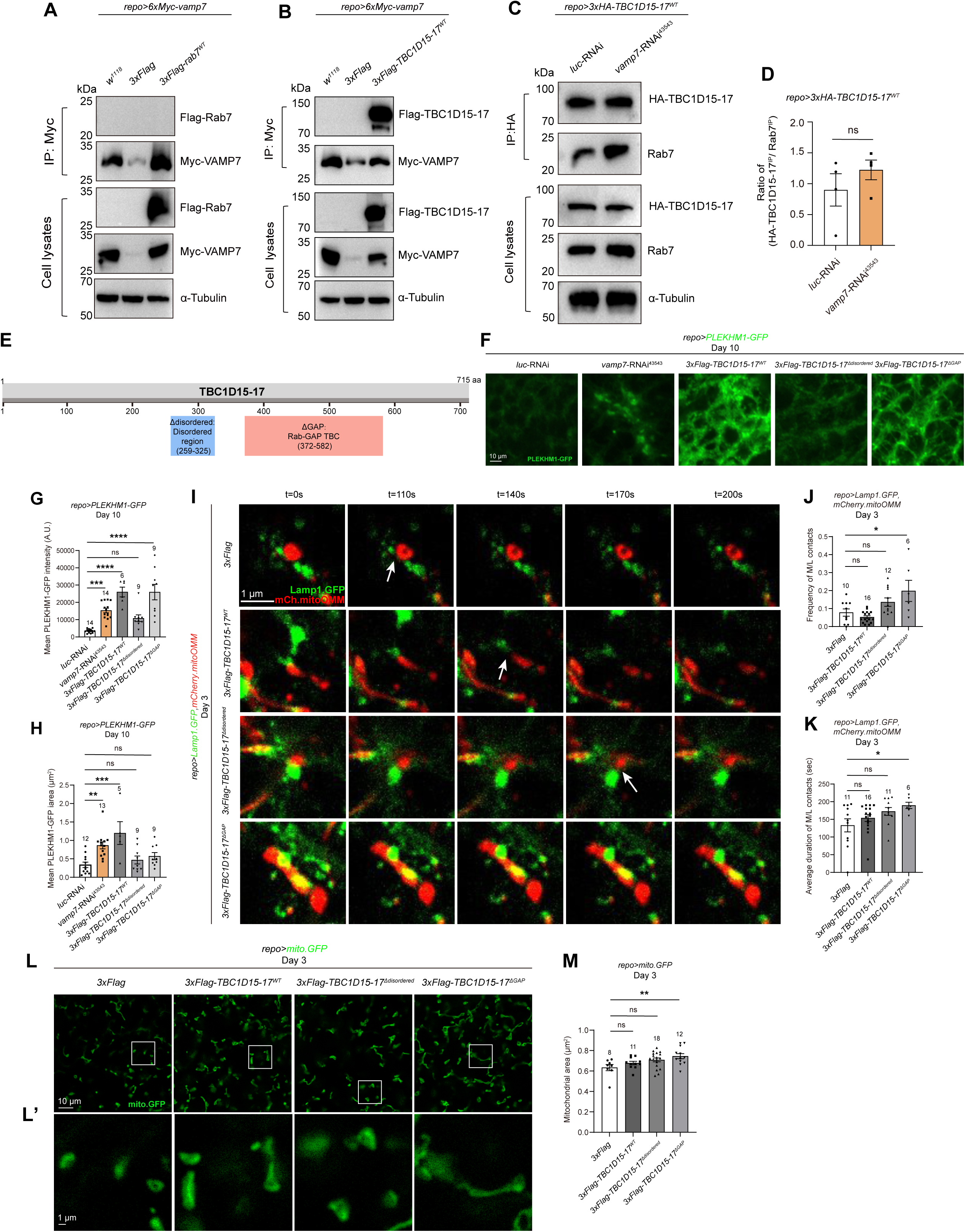
VAMP7 interacts with TBC1D15-17 and expression of TBC1D15-17 lacking the GAP activity causes increased GTP-bound Rab7 levels, impaired MLCs, and mitochondrial elongation in adult fly glia. (**A** and **B**) Representative WB images for Co-IP interaction between VAMP7 and Rab7 (A) or TBC1D15-17 (B). Note that VAMP7 interacts with TBC1D15-17, not Rab7. (**C** and **D**) Representative WB images (B) and quantifications (C) of TBC1D15-17-Rab7 interaction with or without glial VAMP7. Cell lysates from the adult fly heads expressing 3xHA-TBC1D15-17 and *luc-*RNAi or *vamp7-*RNAi^43543^ were collected and immunoprecipitated. Note that TBC1D15-17 remains associated with Rab7 upon glial VAMP7 reduction. (**E**) An illustration of the TBC1D15-17 protein structure. The disordered domain (259-325aa) and GAP domain (372-582aa) are labeled. (**F**-**H**) Representative images (F) and quantifications (G and H) of transgenic flies expressing PLEKHM1(CG6613)-GFP and truncated TBC1D15-17 variants. Note that GFP signals exhibit aggregated patterns in forms of multiple puncta and elevated intensities when expressing *vamp7*-RNAi^43543^, *3xFlag-TBC1D15-17^wt^*, or *3xFlag-TBC1D15-17^ΔGAP^*. (**I**-**M**) Representative images (I and L) and quantifications (J,K,M) on *in-vivo* time lapse analysis of mitochondria-lysosome contacts (MLCs, I) and mitochondrial morphology (L) in adult fly glia. Two reporters: *UAS-mCherry.mitoOMM* (labels the mitochondrial outer membrane, red) and *UAS-Lamp1.GFP* (labels lysosome, green) were expressed in glia to label MLCs, whereas mitochondrial morphology was labeled by *UAS-mito.GFP* (green). Note that TBC1D15-17^ΔGAP^ expression in glia causes prolonged MLCs and elongated mitochondria. The selected white squares in L are enlarged and aligned in L’. All samples examined are brains of 3-day-old adult flies unless noted otherwise. For single transgene experiment, *luc*-RNAi was used as a control. For double transgene experiment, *UAS-LacZ*; *luc*-RNAi was used as a control. White arrows indicate the untethering (I). Scale bars of different sizes are indicated on the images. Serial confocal Z-stack sections were taken at similar planes across all genotypes, showing representative single layer or merged (MLCs) images. Statistical graphs are shown with scatter dots and the number of brain (one MLC per brain) listed on top. Images for the time lapse series are shown in 3x3 μm squares centering on the target MLC selected from the movies (Movies S13-S16). Parameters for the contact dynamics in J and K are quantified using Imaris and ImageJ. For Co-IP analysis, 200 independent fly brains for each genotype were analyzed. Data are shown as mean ± SEM. P-values of significance (indicated with asterisks, ns no significance, * p<0.05, ** p<0.01, *** p<0.001 and **** p<0.0001) are calculated by one-way ANOVA followed by Tukey’s multiple comparisons test.

### Expression of TBC1D15-17 lacking the GAP activity increases GTP-bound Rab7 levels, impairs the untethering of MLCs, and causes mitochondrial elongation in adult fly glia

TBC1D15-17 contains multiple domains including a TBC domain that mediates GAP activity and an uncharacterized disordered domain (Figure 5E). Given that mutations in mammalian TBC1D15 disrupting its GAP activity, like D397A or R400K in the TBC domain, are not conserved in fly TBC1D15-17^15^, transgenic flies expressing the truncated TBC1D15-17 lacking either the entire TBC domain (*UAS-3xFlag-TBC1D15-17^ΔGAP^*) or the uncharacterized disordered domain (*UAS-3xFlag-TBC1D15-17^Δdisordered^*) were generated. Consistent with the *vamp7*-RNAi results, TBC1D15-17^ΔGAP^ expression caused increased intensities and area of PLEKHM1-GFP in adult fly glia, whereas TBC1D15-17^Δdisordered^ failed to do so (Figures 5F-5H). TBC1D15-17 expression also caused an increase in PLEKHM1-GFP levels, suggesting that other regulations exist for TBC1D15-17-mediated Rab7 GTP hydrolysis. Consequently, the dynamics of MLCs were analyzed upon expression of different TBC1D15-17 variants in adult fly glia. Whereas the control MLCs untethered around t=110s, MLCs expressing TBC1D15-17^WT^ or TBC1D15-17^Δdisordered^ untethered at a time scale similar to that of the control, at t=140s or 170s, respectively (white arrows in top three panels, Figure 5I). On the other hand, TBC1D15-17^ΔGAP^ expression significantly increased the frequency and duration of the MLCs (Figures 5I-5K). Consistent with these results, mitochondrial elongation was also detected upon TBC1D15-17^ΔGAP^ expression in glia (Figures 5L and 5M). Taken together, these results suggest that TBC1D15-17 GAP activity contributes to the untethering of MLCs and mitochondrial dynamics in glia.

### TBC1D15-17 GAP activity contributes to VAMP7-mediated MLCs untethering and mitochondrial dynamics in adult fly glia

We next asked if VAMP7 regulates the dynamics of MLCs and glial mitochondrial elongation via TBC1D15-17 function. While the MLCs expressing two control *UAS* transgenes untethered at around t=130s, the MLCs in VAMP7-deficient glia failed to untether within the time frame examined (Figures 6A-6C). Interestingly, the MLCs co-expressing *vamp7*-RNAi^43543^ and TBC1D15-17^WT^ or TBC1D15-17^Δdisordered^ started to untether at a similar time scale to the control at around t=90s and 50s, respectively (white arrows in the middle panel, Figure 6A). The increased frequency and duration of the contacts were significantly suppressed upon co-expression, except that the duration of the contacts co-expressing TBC1D15-17^Δdisordered^ and *vamp7*-RNAi was decreased without significance (Figures 6B and 6C). Notably, TBC1D15-17^ΔGAP^ expression failed to suppress the increase in frequency and duration of the MLCs expressing *vamp7*-RNAi^43543^ (Figures 6A-6C). Furthermore, expression of TBC1D15-17^WT^ or TBC1D15-17^Δdisordered^, but not TBC1D15-17^ΔGAP^, significantly suppressed the mitochondrial elongation upon glial VAMP7 reduction (Figures 6D and 6E). Taken together, these results indicate that VAMP7 regulates the dynamics of MLCs and mitochondrial dynamics via interacting with TBC1D15-17 in adult fly glia, with TBC1D15-17 GAP activity contributing to both events.

**Figure 6.**
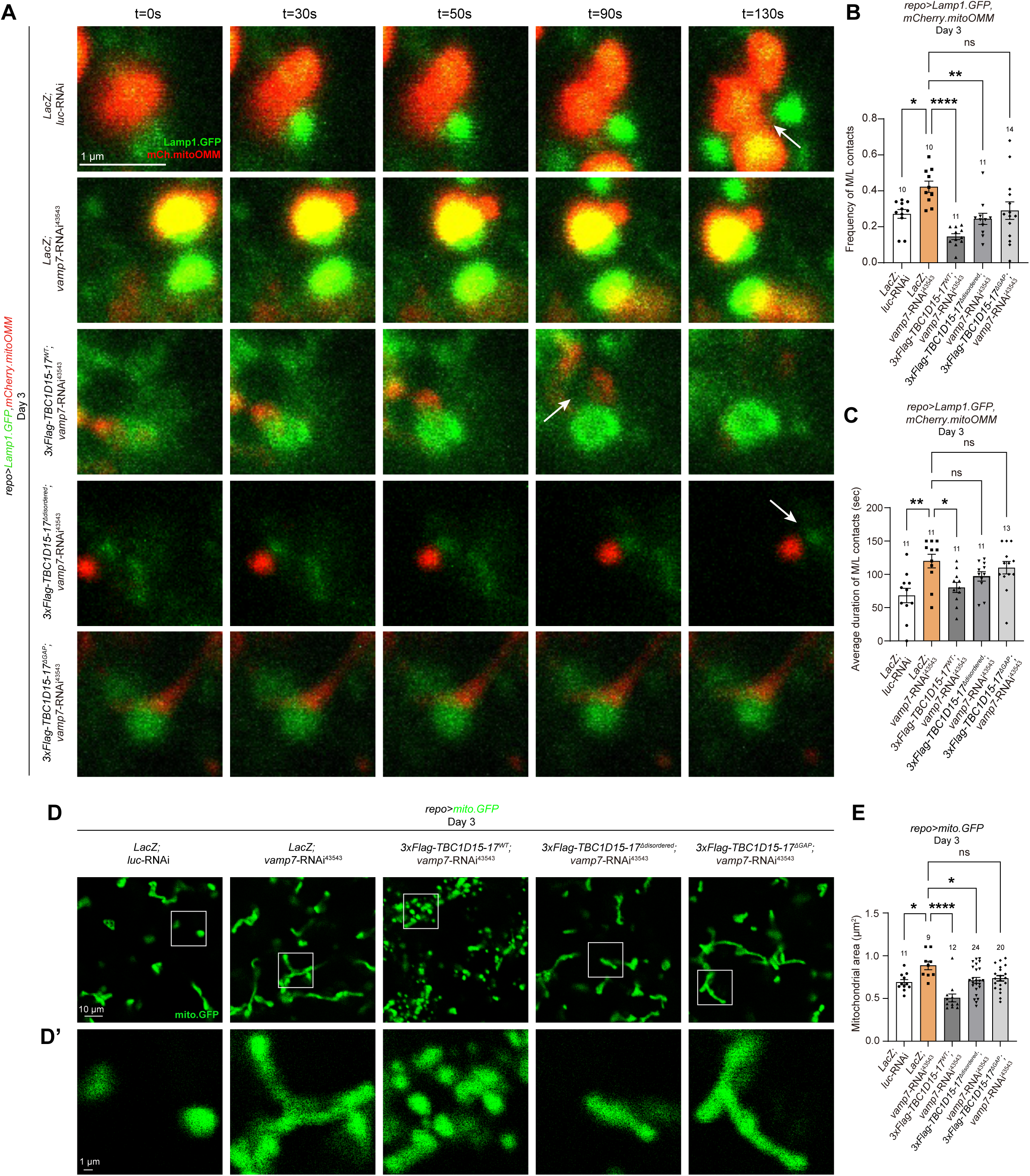
VAMP7 regulates mitochondria-lysosome contacts via Rab7 GAP TBC1D15-17 in adult fly glia. (**A**-**E**) Representative images (A,D) and quantifications (B,C,E) on *in-vivo* time lapse analysis of mitochondria-lysosome contacts (MLCs, A) and mitochondrial morphology (D) in adult fly glia. Two reporters: *UAS-mCherry.mitoOMM* (labels the mitochondrial outer membrane, red) and *UAS-Lamp1.GFP* (labels lysosome, green) were expressed in glia to label MLCs, whereas mitochondrial morphology was labeled by *UAS-mito.GFP* (green). Whereas the contacts persist up to t=130s upon glial VAMP7 reduction, co-expression of TBC1D15-17^WT^ or TBC1D15-17^Δdisordered^ restores the prolonged contact dynamics (t=90s and 50s, middle two panels in A). Note that TBC1D15-17^ΔGAP^ expression fails to restore the frequency and the duration of the contacts (B and C). The average mitochondrial area quantified by ImageJ is significantly restored when co-expressing *vamp7*-RNAi^43543^ and TBC1D15-17^WT^ or TBC1D15-17^Δdisordered^, but not TBC1D15-17^ΔGAP^ (D and E). The selected white squares in D are enlarged and aligned in D’. All samples examined are brains of 3-day-old adult flies unless noted otherwise. For single transgene experiment, *luc*-RNAi was used as a control. For double transgene experiment, *UAS-LacZ*; *luc*-RNAi was used as a control. White arrows indicate the untethering (A). Scale bars of different sizes are indicated on the images. Serial confocal Z-stack sections were taken at similar planes across all genotypes, showing representative single layer (mitochondrial morphology) or merged (MLCs) images. Statistical graphs are shown with scatter dots and the number of brains (one MLC per brain) listed on top. Images for the time lapse series are shown in 1.82x1.82 μm squares centering on the target MLC selected from the movies (Movies S17-S21). Parameters for the contact dynamics in B and C are quantified using Imaris and ImageJ. Data are shown as mean ± SEM. P-values of significance (indicated with asterisks, ns no significance, * p<0.05, ** p<0.01, *** p<0.001 and **** p<0.0001) are calculated by one-way ANOVA followed by Tukey’s multiple comparisons test.

### Reducing *vamp7* expression causes mitochondrial dysfunction in adult fly glia

To further demonstrate that VAMP7 regulation of MLCs and mitochondrial dynamics in glia are functionally relevant, we assessed the levels of various reporters indicative of mitochondrial function. As mitochondria are the major sites for ROS production, control and *repo>vamp7-*RNAi^43543^ adult fly brains were examined for ROS levels with the dihydroethidium (DHE) staining. Consistent with previous findings, DHE-positive fluorescent intensities increased when downregulating the expression of superoxide dismutase 1 (SOD1), the enzyme that regulates ROS levels by catalyzing the conversion of superoxide into oxygen and hydrogen peroxide (Figure 7A). Interestingly, *vamp7-*RNAi^43543^ expression in glia caused a similar increase, suggesting that glial VAMP7 regulates cellular ROS levels (Figure 7A). In addition, the transcription levels of genes related to the mitochondrial fatty acid β-oxidation pathway, including those encoding acyl-CoA dehydrogenases (*CG17544* and *CG9527*) and fatty acid synthase (*FASN*), were increased in adult fly brains expressing *vamp7*-RNAi^43543^ in glia^28, 29^ (Figure 7B). Notably, the levels of BODIPY-positive LDs increased in both larval and adult fly brains upon glial VAMP7 reduction (Figures 7C-7E), with a similar increase in the overall LD triacylglycerol (TAG) content (Figure 7F). The increased LD number was reduced upon adding the potent antioxidant α-Lipoic acid in 3-day-old adult fly brains (Figures 7G and 7H). Taken together, these results indicate that glial VAMP7 regulates mitochondrial function, ROS levels, and LD production in the brain.

**Figure 7.**
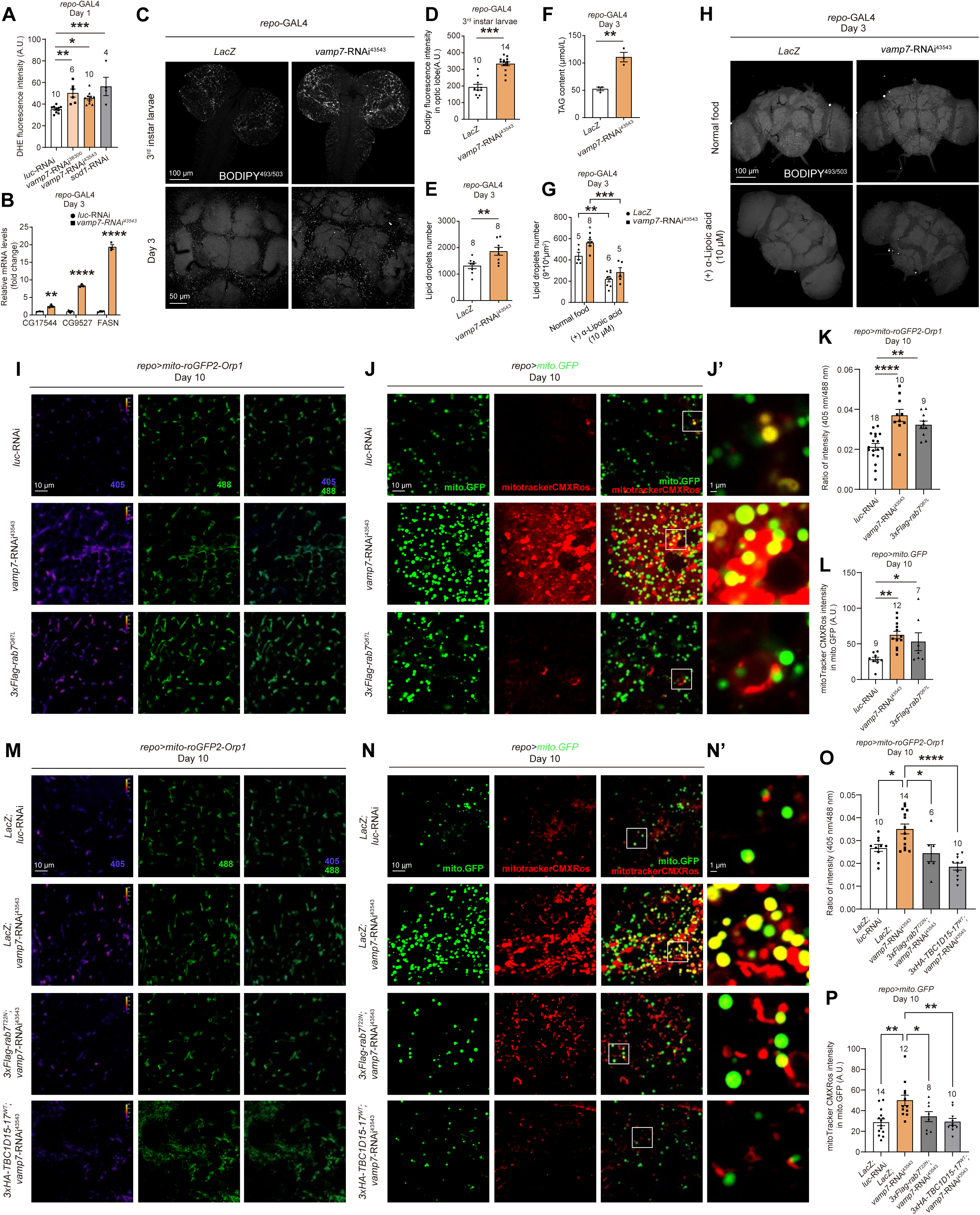
Silencing *vamp7* expression increases ROS levels and disrupts mitochondrial function in adult fly glia. (**A**) DHE fluorescent intensities indicating cellular ROS levels are significantly increased upon glial VAMP7 or SOD1 reduction. (**B**) RT-qPCR analyses reveal increased transcription levels of genes related to mitochondrial function including the ones encoding acyl-coA dehydrogenases (*CG17544* and *CG9527*) and fatty acid synthase (*FASN*). (**C**-**H**) The intensity and number of LDs labeled by the BODIPY dye in both larval and adult fly brains (C-E) and the TAG content in the adult fly brains (F) are significantly increased upon glial VAMP7 reduction. Note that the intensities of LDs are quantified in selected regions of 3^rd^ instar larval brains (D) vs. the whole brains in 3-day-old adult flies (E). Glial VAMP7 regulates LD formation as the addition of α-lipoic acid suppresses ROS levels in both control and *repo>vamp7-*RNAi^43543^ adult fly brains (G and H). (**I-P**) Representative images (I and M) and quantifications (K and O) of the mitochondrial H_2_O_2_ reduction-oxidation activity monitored by the reporter *UAS-mito-roGFP-Orp1* in 10-day-old *repo>mito-roGFP-Orp1* adult fly brains. Note that the mitochondrial H_2_O_2_ reduction-oxidation activity is upregulated when expressing *vamp7*-RNAi^43543^ or Rab7^Q67L^ (I and K). The increased activity upon glial VAMP7 reduction is suppressed by co-expressing Rab7^T22N^ or TBC1D15-17^WT^ (M and O). Representative images (J,J’,N,N’) and quantifications (L and P) of the MitoTracker Red CMXRos fluorescent intensities detecting cellular mitochondrial membrane potential per mito.GFP-positive mitochondria. Note that the mitochondrial membrane potential is hyperpolarized when expressing *vamp7*-RNAi^43543^ or Rab7^Q67L^ (J,J’,L). The hyperpolarization in VAMP7-deficient glia is suppressed by co-expressing Rab7^T22N^ or TBC1D15-17^WT^ (N,N’,P). The selected white squares in J and N are enlarged and aligned in J’ and N’. All samples examined are brains of 3-day-old adult flies unless noted otherwise. For single transgene experiment, *UAS-LacZ* (*LacZ*) or *luc*-RNAi was used as a control. For double transgene experiment, *UAS-LacZ*; *luc*-RNAi was used as a control. Scale bars of different sizes are indicated on the images. Serial confocal Z-stack sections were taken at similar planes across all genotypes, showing representative single layer or merged (LD) images. Statistical graphs are shown with scatter dots and the number of brains listed on top. For TAG statistics, 20 independent fly brains for each genotype were analyzed. Data are shown as mean ± SEM. P-values of significance (indicated with asterisks, ns no significance, * p<0.05, ** p<0.01, and *** p<0.001) are calculated by two-tailed unpaired t-test compared two groups experiments or by one-way ANOVA compared three groups followed by Tukey’s multiple comparisons test.

We also investigated mitochondrial function by analyzing the activity of the reporter *UAS-mito-roGFP2-Orp1*^30^. Interestingly, the 405/408 intensities reflecting the H_2_O_2_ reduction-oxidation activity in the mitochondrial matrix were significantly increased in 10-day-old adult fly glia expressing Rab7^Q67L^ or *vamp7*-RNAi^43543^ (Figures 7I and 7K). Staining the brains with the dye mitotrackerCMXROS, indicative of the living mitochondria membrane potential, also revealed increased intensities upon Rab7^Q67L^ or *vamp7*-RNAi^43543^ expression in glia (Figures 7J and 7L). Results from these quantitative analyses indicate that VAMP7 regulates mitochondrial function in glia. Importantly, both the 405/408 intensities and the mitotracker dye levels were suppressed when co-expressing the GDP-bound Rab7^T22N^ or TBC1D15-17^WT^ in glia (Figures 7M-7P), suggesting that altering Rab7 GTP hydrolysis by increasing GDP-bound Rab7 or Rab7 GAP activity in VAMP7-deficient glia reestablishes the balance of Rab7 GTP/GDP levels, hence recovering the mitochondrial function in adult fly glia.

### Glial VAMP7 contributes to DA neuron survival

As glia-neuron crosstalk underlies neuronal function and survival, we next investigated if VAMP7-mediated mitochondrial dynamics in glia contribute to DA neuron survival. DA neurons are located in clusters and named according to their relative positions in adult fly brains^31, 32^. These clusters include protocerebral posterior lateral (PPL)1, PPL2, protocerebral posterior medial (PPM)1, PPM2, and PPM3 (Figure 8A). Interestingly, silencing glial *vamp7* expression caused DA neuron loss at PPM1/2, PPM3, and PPL1 clusters in an age-dependent manner (white arrows, Figures 8B-8D). In addition, *vamp7*-RNAi^43543^ expression in glia in the presence of *Elav*-GAL80, which inhibits GAL4 expression in neurons, also caused DA neuron loss in the PPM1/2 cluster, further demonstrating that DA neuron loss is a result explicitly of glial VAMP7 knockdown (Figures 8E and 8F). Given that DA neurodegeneration in the PPM1/2 cluster has been implicated in a *Drosophila* PD model overexpressing α-synuclein^33^, it is conceivable that glial VAMP7 plays important roles in regulating DA neuron survival under pathological conditions.

**Figure 8.**
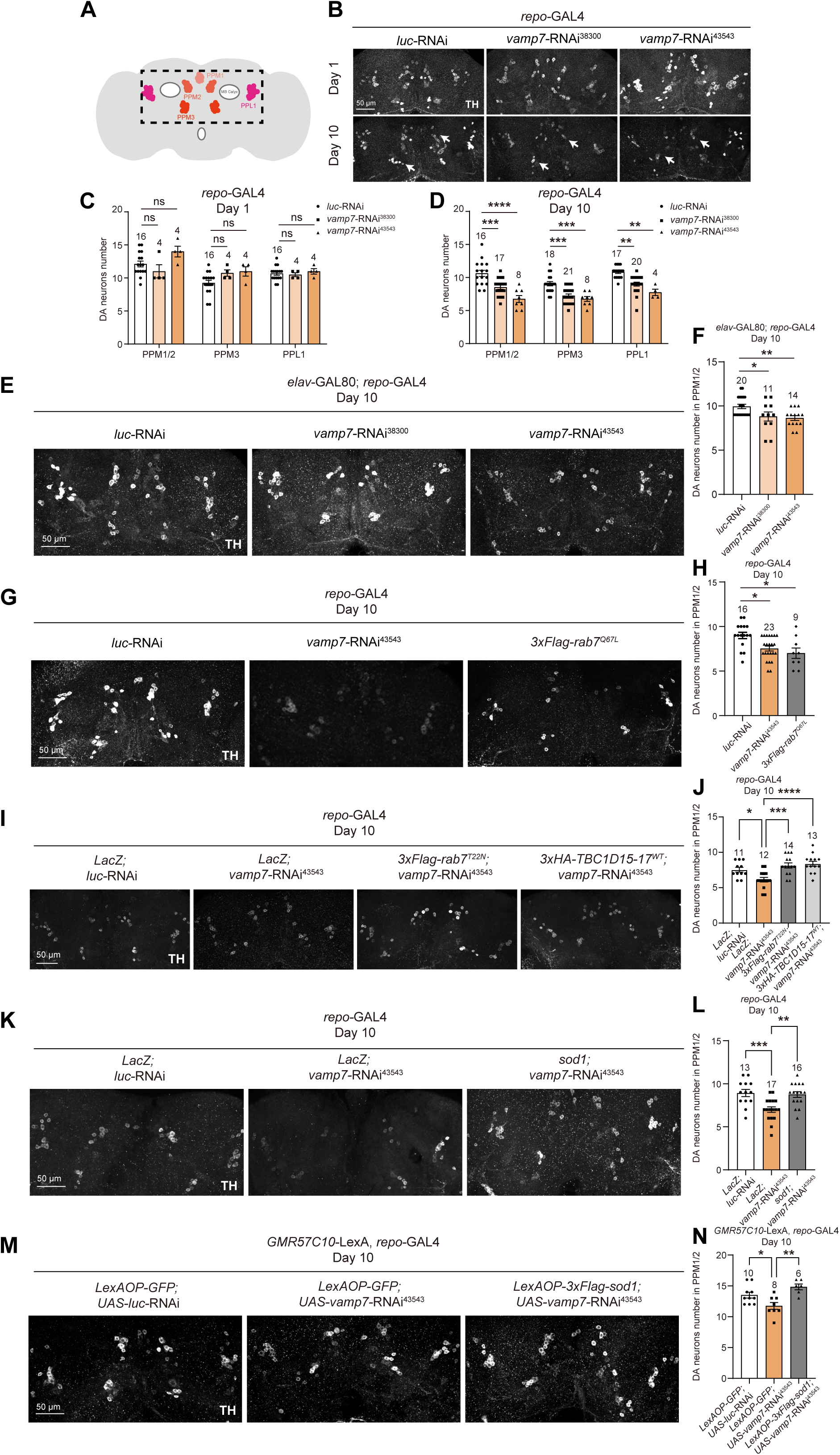
Glial VAMP7 contributes to DA neuron survival. (**A**) An schematic diagram illustrating the DA neuron clusters in adult fly brains. The black dashed box encloses the area containing DA neuron clusters (PPM1/2, PPL1, and PPM3) and is visualized in B,E,G,I,K,M. (**B**-**D**) The numbers of DA neurons in PPM1/2, PPM3, and PPL1 are significantly decreased (white arrows in B) in the aged flies (10-day-old) upon glial VAMP7 reduction. (**E-N**) Representative images (E,G,I,K,M) and quantifications (F,H,J,L,N) of DA neuron number in the PPM1/2 cluster under different genetic conditions. Note that DA neuron number is consistently decreased in the 10-day-old flies expressing *vamp7*-RNAi^43543^ in the presence of Elav-Gal80, which blocks Gal4 expression in neurons (E and F). DA neuron number is also significantly decreased in the 10-day-old flies expressing Rab7^Q67L^ (G and H). The DA neurodegeneration upon glial VAMP7 reduction is restored by co-expressing Rab7^T22N^ or TBC1D15-17^WT^ (I and J). Expressing SOD1 in glia (K and L) or neurons (M and N) restores the DA neuron loss in PPM1/2 upon glial VAMP7 reduction. DA neurons are detected with anti-TH antibodies. All samples examined are brains of 10-day-old adult flies unless noted otherwise. For single transgene experiment, *luc*-RNAi was used as a control. For double transgene experiment, *UAS-LacZ*; *luc*-RNAi was used as a control. Scale bars of different sizes are indicated on the images. Serial confocal Z-stack sections were taken at similar planes across all genotypes, showing representative merged images. Statistical graphs are shown with scatter dots and the number of brains listed on top. Data are shown as mean ± SEM. P-values of significance (indicated with asterisks, ns no significance, * p<0.05, ** p<0.01, *** p<0.001 and **** p<0.0001) are calculated by ordinary one-way ANOVA followed by Tukey’s multiple comparisons test.

On the other hand, whereas Rab7^Q67L^ expression in glia caused similar DA neuron loss in the PPM1/2 cluster (Figures 8G and 8H), co-expression of Rab7^T22N^ or TBC1D15-17 with *vamp7*-RNAi^43543^ suppressed the degeneration, suggesting that reestablishing the balance of Rab7 GTP/GDP levels, potentially restoring the dynamics of MLCs in glia, promotes DA neuron survival (Figures 8I and 8J). Finally, suppressing the ROS levels by expressing SOD1 in either glia or neurons restored the DA neurodegeneration in *vamp7*-RNAi^43543^-expressing brains, suggesting that modulating ROS levels in either glia or neurons promotes DA neuron survival (Figures 8K-8N). These results further implicate a role of glial mitochondrial dynamics in DA neuron survival, as tuning down the ROS levels in glia only also helps to ameliorate DA neurodegeneration.

## Discussion

SNARE proteins are canonical regulators of membrane fusion. Our findings uncover a new role for the SNARE protein VAMP7 in glial mitochondrial dynamics. Silencing *vamp7* expression in glia causes fission/fusion-dependent mitochondrial elongation, owing to the dysregulated untethering of MLCs. VAMP7 interacts with the Rab7 GAP TBC1D15-17 and regulates Rab7 GTP/GDP levels, hence modulating the dynamics of MLCs. Mitochondria in VAMP7-deficient glia are dysfunctional with hyperpolarized membrane potential, leading to increased ROS activity and lipid droplet formation in the brain. These deficits endanger nearby DA neurons, linking glial mitochondrial dynamics to DA neuron survival (Figure 9).

**Figure 9.**
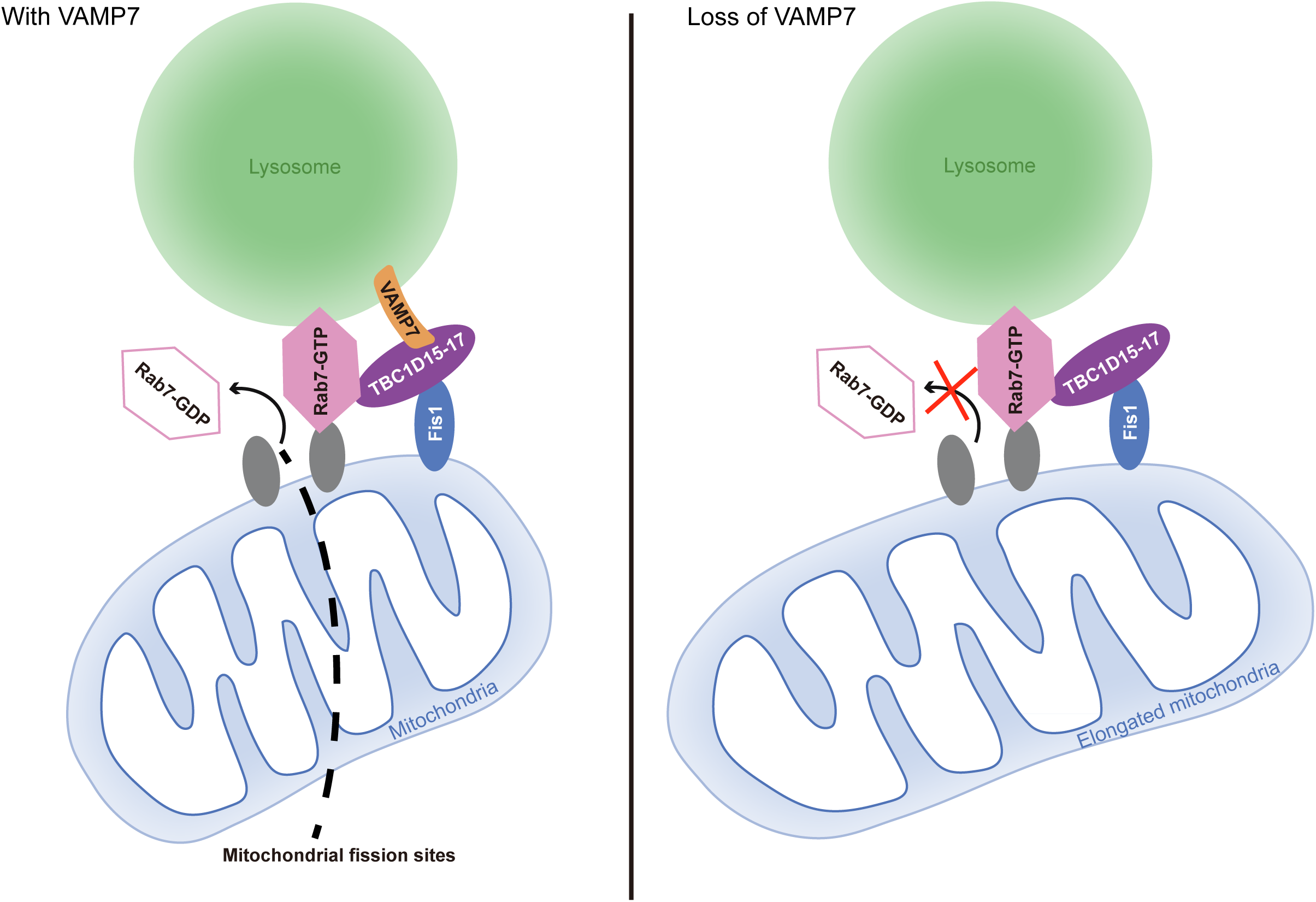
VAMP7 regulates mitochondria-lysosome contacts in glia. An illustrated mechanism of VAMP7-mediated mitochondrial dynamics in glia. Under physiological conditions, VAMP7 interacts with TBC1D15-17, facilitating Rab7 GTP hydrolysis for the untethering of MLCs in adult fly glia. The contacts determine the site for fission. Upon glial VAMP7 reduction, TBC1D15-17 remains associated with Rab7, yet is unable to promote Rab7 GTP hydrolysis, leading to persistent contact and mitochondrial elongation.

### VAMP7 mediates mitochondrial dynamics in glia

As the untethering of MLCs depends on Rab7 GTP hydrolysis, we uncovered that the persistent contact (i.e., dysregulated untethering) induced by decreased *vamp7* expression in glia is linked to a disrupted balance of Rab7 GTP/GDP levels; Rab7 GTP levels are significantly increased upon silencing glial *vamp7* expression. VAMP7 interacts with the Rab7 GAP TBC1D15-17 to promote Rab7 GTP hydrolysis. Without VAMP7, TBC1D15-17 remains associated with Rab7, yet the TBC1D15-17 GAP activity is likely impaired given the increased GTP-bound Rab7 levels. Expression of TBC1D15-17 lacking the GAP activity, like *vamp7*-RNAi, also causes prolonged MLCs and mitochondrial elongation in adult fly glia. These results suggest that VAMP7 interaction with TBC1D15-17 promotes Rab7 GTP hydrolysis, hence regulating the MLCs untethering and mitochondrial dynamics in glia.

### VAMP7 mediates glial mitochondrial dynamics independently of autophagosome-lysosome fusion and mitophagy

Consistent to a role of VAMP7 in autophagosome-lysosome fusion, the autophagic flux we monitored using a dual reporter *UAS-GFP-mCherry-Atg8a* when expressing *vamp7*-RNAi in glia was significantly impaired (Figures S2A and S2B). The Pearson’s coefficient values for colocalization of GFP and mCherry-positive signals were significantly increased upon glial VAMP7 reduction, indicating non-quenched GFP signals due to impaired fusion (Figure S2B). Nonetheless, the glial mitochondrial area was not altered in the presence of Bafilomycin A1 (BafA1), a drug blocking autophagosome-lysosome fusion (Figures S2C and S2D)^34, 35, 36^. Despite that the STX17-SNAP29-VAMP7 complex regulates autophagosome-lysosome fusion, STX17 inhibition in glia failed to induce mitochondrial elongation (Figures S2E and S2F). DA neuron number also remained unaffected upon STX17 or SNAP29 inhibition in glia (Figures S2G and S2H). Thus, it is likely that VAMP7 regulates the untethering of MLCs independently of its function in the autophagosome-lysosome fusion.

Consistent with the results of disrupting autophagosome-lysosome fusion, inhibiting autophagy by downregulating the expression of *atg1* or *atg8* in glia causes smaller mitochondria, suggesting enhanced mitochondrial fission (Figures S2I and S2J)^37, 38^. Given that *vamp7*-RNAi expression induces mitochondrial elongation in glia, these results reinforce a role for VAMP7 in glial mitochondrial dynamics independent of autophagy. On the other hand, whereas the colocalization of Lamp1.GFP- and mitoOMM.mCherry-positive puncta is significantly increased (as shown throughout the study as prolonged contacts, Figure S2K, S2L, and S2N). Atg8a was very minimally localized to the MLCs upon glial VAMP7 reduction (Figures S2K, S2M, and S2O). These results are consistent with the notion that elongated mitochondria are resistant to and slow down mitophagy; reduced glial *vamp7* expression increases the formation of MLCs yet inhibits mitophagy.

### VAMP7-mediated mitochondrial dynamics in glia functionally contribute to DA neuron survival

Our findings further support the hypothesis that MLCs mark the sites for mitochondrial fission. Upon reduction in glial *vamp7* expression, the untethering of MLCs is hindered, disrupting subsequent fission, leading to elongated mitochondria, which is suppressed by the expression of the fission/fusion factors. Furthermore, these elongated mitochondria are dysfunctional with hyperpolarized membrane potential, enhanced ROS activity, and lipid droplet formation in the brain. These changes are at the whole-brain level and affect DA neuron survival. Reducing the ROS activity by expressing SOD1 in glia or neurons suppresses the DA neurodegeneration, supporting the notion that VAMP7-mediated mitochondrial dynamics underlie the mechanism of glia-neuron crosstalk contributing to neuron survival.

### SNARE proteins in glial mitochondrial dynamics

To date, no evidence indicates that SNARE proteins mediate mitochondrial fusion. A recent report reveals that the STX17-SNAP29-VAMP7 complex contributes to the megamitochondria engulfing lysosome (MMEL) under hypoxia^36^. STX17 has also been reported to regulate the activity of the fission factor Drp1^39, 40^. Notably, STX17 localizes to the ER-mitochondria contacts to execute its function in mitochondrial fission^39, 41^, while we uncover a role for VAMP7 in the untethering of MLCs. Based on these findings, it is feasible to speculate that SNARE proteins are significant players in the contacts of different organelles, in addition to their function in membrane fusion.

Similar to neurons, glia, particularly astrocytes, possess SNARE machineries for their vesicular trafficking function. These SNARE machineries, some in the same composition as in neurons and some different, are responsible for slow secretory exocytosis rather than fast and immediate release of signaling molecules^42^. Very importantly, they regulate synaptic transmission and plasticity, antigen presentation in neuro-inflammation, and control different types of behavior. Our findings unexpectedly reveal a new role for the canonical SNARE protein VAMP7 in glial mitochondrial dynamics. Considering that mitochondria are present in astrocytic processes in close proximity to neurons and contribute to local calcium signaling, ATP production, and metabolism of signaling molecules, it is conceivable that VAMP7-mediated glial mitochondrial dynamics participate in the metabolic process supporting neuron survival. In support of the hypothesis, downregulating *vamp7* expression in the fly astrocyte-like glia also caused DA neuron loss (Figure S3). Given that DA neurodegeneration is one of the hallmarks in PD, our findings illustrate a SNARE-dependent mechanism controlling organelle contacts, glial mitochondrial dynamics, and DA neuron survival. Canonical SNARE proteins not only regulate membrane fusion, but also are important mediators in mitochondrial dynamics; future work is warranted to explore additional roles of SNARE proteins in various cellular contexts in both cell-autonomous and non-cell-autonomous fashions.

## Acknowledgments

We thank Bloomington *Drosophila* Stock Center, Vienna *Drosophila* RNAi Center, Tsinghua Fly Center, the Core Facility of *Drosophila* Resource and Technology, Shanghai Institute of Biochemistry and Cell Biology, Chinese Academy of Sciences, Developmental Studies Hybridoma Bank, Chao Tong, Chun-Hong Chen, Henry Sun, and Aike Guo for fly stocks and antibodies. We also thank the Molecular Imaging Core Facility (MICF), the Molecular and Cell Biology Core Facility (MCBCF), and the Multi-Omics Core Facility (MOCF) at the School of Life Science and Technology, ShanghaiTech University for providing technical support; Shengxiong Wang in Shanghai JiaoTong University School of Medicine and Yu Kong and Lijun Pan in Electron Microscopy Facilities of Center for Excellence in Brain Science and Technology, Chinese Academy of Science for assistance with TEM sample preparation; The Electron Microscopy Center in the School of Physical Science and Technology, ShanghaiTech University for providing TEM technical support; DroBot Biotechnology for quality fly food supply, delicate fly-keeping service, and experimental device design; Ho lab members and Yan Zou for discussion and comments. This work was supported by intramural grants from National Yang Ming Chiao Tung University, National Natural Science Foundation of China (32170962 to M.S.H, 32371067 to Y.P), Shenzhen Medical Academy of Research and Translation (B2402005 to Y.P.), 2030 Cross-Generation International Outstanding Young Scholars Program National Science and Technology Council Taiwan (113-2628-B-A49-003), and Brain Research Center National Yang Ming Chiao Tung University from The Featured Areas Research Center Program within the framework of the Higher Education Sprout Project by the Ministry of Education (MOE) in Taiwan.

## Author Contributions

H.W and M.S.H conceived and designed the study. H.W, M.W, C.L, Y.L, L.W, S.Y, and S.Z performed the experiments. H.W, Y.T, Y.C, Y.L, Y.P, and M.S.H analyzed the data. M.S.H wrote the paper with input from H.W and Y.P. All authors read and approved the manuscript.

## Materials and Methods

### Drosophila genetics

Flies were raised under standard conditions at 25[with 70% humidity. All fly crosses were carried out at 25°C with standard laboratory conditions unless noted otherwise. All strains were obtained from Bloomington *Drosophila* Stock Center (BDSC), the Vienna *Drosophila* RNAi Center (VDRC), Tsinghua Fly Center, or as gifts from colleagues. The following fly strains were used in this study: *repo-GAL4* (a pan-glial driver), *UAS-luciferase-IR* (BL31603), *UAS-LacZ* (BL1777), *UAS-mito.HA.GFP* (BL8442), *UAS-mCherry.mitoOMM* (BL66532)*, UAS-Lamp1.GFP* ^43^, *UAS-vamp7-RNAi* (BL38300 and BL43543), *UAS-marf RNAi* (BL31157), *UAS-opa1 RNAi* (THU0811), *UAS-sod1 RNAi* (THU3207), *UAS-mito-roGFP2-Orp1* (BL67667), *UASp-GFP-mCherry-Atg8a* (Henry Sun), *UAS-HRP* (Aike Guo), *UAS-drp1* (Chao Tong), and transgenic flies made in the present study (*UAS-6xMyc-vamp7, UAS-3xFlag-Rab7^wt^*, *UAS-3xFlag-Rab7^Q67L^, UAS-3xFlag-Rab7^T22N^, UAS-3xHA-TBC1D15-17, UAS-3xHA-RILPL*, *LexAOP-3xFlag-sod1, UAS-TBC1D15-17^wt^, UAS-3xFlag-TBC1D15-17^Δdisordered^, UAS-3xFlag-TBC1D15-17^ΔGAP^*).

### Molecular biology

#### Plasmid cloning

Plasmids were constructed by the polymerase chain reaction (PCR) using DNA templates from *Drosophila*. These plasmids express wild type Rab7 (621 bp), Rab7 point mutant variants including Rab7^Q67L^ (621 bp, Glutamine 67 to Leucine), and Rab7^T22N^ (621 bp, Threonine 22 to Asparagine), VAMP7, TBC1D15-17, RILPL, and PLEKHM1. The corresponding DNA sequences were subcloned into pUAST-attB vector containing a 3xFlag, 3xHA, or GFP epitope tag by GenScript (Nanjing, China). *sod1* gene was subcloned into the LexAOP vector containing a 3xFlag epitope tag by PCR. Primers used for amplification:

Sod1-F: CGACGACAAACTCGAGGTGGTTAAAGCTGTCTGCGET

Sod1-R: ACAAAGATCCTCTAGATTAGACCTTGGCAATGCCAATAACG

Fly microinjection was conducted by the *Drosophila* Core Facility, Institute of Biochemistry and Cell Biology, Chinese Academy of Sciences.

<u>RT-qPCR</u>: Total RNAs were extracted from adult fly heads using TransZol Up (Cat. #ET111-01, TransGen, Beijing, China). cDNAs were reversely transcribed using HiScript III RT SuperMix (Cat. #R323-01, Vazyme, Nanjing, China). RT-qPCR reactions were performed using ChamQ Universal SYBR qPCR Master Mix (Cat. #Q711-02, Vazyme) and ABI 7500 RT-PCR system. The mRNA levels were normalized to *rp49*. The expression levels were analyzed by ΔΔCT method. Primers used for amplification:

*rp49*-F: CCACCAGTCGGATCGATATGC

*rp49*-R: CTCTTGAGAACGCAGGCGACC

*vamp7*-F: GGCTCGCAATTGGATAGTAGT

*vamp7*-R: TGCTTATCGGGGAATGAGCC

*CG17544*-F: *GTACTATGCTCTGACGCGCT*

*CG17544*-R: *GGTGGTCATGTGCTTCTCCA*

*CG9527*-F*: CTGCTTCCCCTGCTGAAGAA*

*CG9527*-R*: GACGTTTCCATCGCTCATGC*

*FASN*-F: *CGCTCCACTCCAAGAACTCG*

*FASN*-R*: CAGGTTTAGTTGTAGGGGCTAGA*

### FACS sorting of adult fly glia

Approximately 100 *repo>mCD8.GFP* adult fly brains were dissected in fresh Schneider’s Insect Medium (Invitrogen, Carlsbad, CA, USA, Cat# 21720024) and transferred them to the phosphate-buffered saline (PBS) dissociation buffer which contains papain (0.18 U/mL; Worthington, Freehold, NJ, USA, Cat# LK003178) and liberase TM (2.5 mg/mL; Roche, Basel, Switzerland, Cat# 5401119001) to dissociate. Note that papain requires pre-activation at 37℃ for 15 minutes prior to use. Ice-cold Schneider’s Insect Medium was used to inactivate enzymes. Samples were filtrated by 40 μm cell strainers (BD Falcon, Franklin Lakes, NJ, USA, Cat# 352340) into flow cytometry tubes (BD Falcon, Cat# 352003) then sorted by the flow cytometer (FACSMelody, BD Biosciences, Bergen County, NJ, USA).

### Immunohistochemistry

Adult fly brains were dissected and fixed in 4% formaldehyde for 45 minutes, then washed with PBT (PBS + 0.3% TX-100) for 3 times and dissected further to remove additional debris in PBS solution. Clean and fixed brains were blocked in PBT solution with 5% Normal Donkey Serum (NDS) or 10% Bovine Serum Albumin (BSA) and subsequently stained with primary antibodies at 4[overnight and secondary antibodies at room temperature for 2 hours. Primary antibodies used: mouse anti-Atg8a (1:1000, gift from Chao Tong) and rabbit anti-TH (1:1000, Cat. #NB300-109, Novus). For all samples, secondary antibodies used were from Jackson ImmunoResearch (West Grove, PA, USA): donkey anti-rabbit Cy3 (1:1000, Cat. #711-166-152), donkey anti-mouse Cy5 (1:500, Cat. #715-175-150). All samples were mounted in Vectashield mounting media (Cat. #H-1500-10, Vector Laboratories, Burlingame, CA, USA).

### *In-vivo* time lapse analysis

Live adult fly brains were dissected and kept in PBS solution. All samples were mounted in PBS, and immediately scanned anteriorly to posteriorly (top to bottom) in a serial Z-stack of average 5 sections, each of 0.125 μm thickness using Nikon C2 confocal microscope (Tokyo, Japan) with a 63X oil objective in the interval of 10 seconds for 5 minutes. Samples in Figure 6A were scanned in a serial Z-stack of 5 sections, each of 0.395 μm thickness using Zeiss LSM 700 (Oberkochen, Germany) with a 63X oil objective in the interval of 10 seconds for 5 minutes. The frequency of Lamp1.GFP and mCherry.mitoOMM contacts was analyzed by Imaris using autoregressive motion tracking mode. Duration of contacts were analyzed by ImageJ.

### Live fluorescent dye imaging

Live adult fly brains were dissected and kept in PBS solution. For the analysis of LDs, brains were incubated with 1 μg/mL BODIPY™ 493/503 (Cat. #D3922, Invitrogen) for 5 minutes at 25°C and/or 10 μM α-Lipoic acid (Cat #D118666, Aladdin) for 3 days. For the analysis of mitochondria-derived ROS levels, brains were incubated with 50 μM DHE (Cat. #D11347, Invitrogen) for 5 minutes at 25°C. For the analysis of mitochondrial membrane potential, brains were incubated with 500 nM MitoTracker RedCMXRos (Cat. #M7512, Invitrogen) solution in PBS for 15 minutes at 25°C. Images were acquired with the Nikon C2 or A1 confocal microscope (Tokyo, Japan) with a 20X or 60X oil objective.

### Transmission electron microscopy

Adult fly brains were fixed, embedded, stained, and dehydrated according to previous protocols^44^. Ultrathin sections (70 nm) of each brain were cut with a Diatome diamond knife on a Leica EM UC7 ultra-microtome (Leica Microsystems, Germany) and collected on copper grids. Images were acquired with Talos L120C transmission electron microscope (Thermo Fisher Scientific, Waltham, MA, UAS) operating at an acceleration voltage of 120 kV.

### Biochemistry

#### Western blot analysis

Adult fly heads were homogenized using a motorized pestle (Cat. #116005500, MP Biomedicals, Irvine, CA, USA) in lysis buffer (0.4% NP-40, 0.2 mM EDTA, 150 mM NaCl, 20% glycerol, 100 mM Tris-HCl pH7.5, 2% Tween 20, 0.5 mM phosphodiesterase inhibitors, and 1 mM PMSF). Proteins were separated on SDS-PAGE gels and transferred to PVDF membranes (Cat. #IPFL00010, Millipore, Billerica, MA, USA). The membranes were blocked by 5% fat-free milk diluted in PBS with 0.1% Tween-20 for 30 minutes. Samples were incubated with the primary antibodies at 4°C overnight and HRP-conjugated secondary antibodies at room temperature for 2 hours. Primary antibodies used: mouse-anti-HA (1:1000, Cat. #12C10, Roche), rabbit anti-Myc (1:2000, Cat. #0912-2, Hua An Biotechnology), mouse anti-Rab7 (1:100, DSHB), rabbit-anti-α-Tubulin (1:5000, Cat. #SB-AB0049, Share Biotechnology). Bands were visualized by Clarity Western ECL Substrate (Cat. #WBKLS0500, Millipore). Results are representative of at least three biological replicates.

### Co-immunoprecipitation

The cell lysates were collected as described above for centrifugation at 12,000 rpm, at 4°C for 30 minutes, the soluble supernatant was incubated with prewashed anti-HA magnetic beads (Cat. # HY-K0201; MedChemExpress), anti-c-Myc magnetic beads (Cat.# HY-K0206A, MedChemExpress) at 4°C overnight. After pull-downs, gels were extensively washed in lysis buffer for 30 minutes 3 times and then subjected to western blot analysis. The following antibodies were used for blotting: rabbit-anti-Myc (1:2000, Cat. #0912-2, Hua An Biotechnology), mouse-anti-Flag (1:1000, Cat. #F3165, Sigma), mouse-anti-HA (1:1000, Cat. #3F10, Roche), mouse anti-Rab7 (1:100, DSHB) and rabbit-anti-α-Tubulin (1:5000, Cat. #SB-AB0049, Share Biotechnology).

### BafilomycinA1 treatment

BafA1 (Cat. #BML-CM110-0100, ENZO) was diluted in DMSO (Cat. #D8371-50, solarbio) in a concentration of 1 mM, then added to 0.5 ml of liquid yeast for a final concentration of 4 μM. Yeast containing BafA1 was added to the top of the fly food in standard vials kept overnight to allow the evaporation of residual DMSO. Flies of different genotypes (20 females and 20 males) were then put in each vial and kept at 25°C for 5 days. After treatment, flies were collected for imaging analysis. The TAG content in the LD was quantified by an triglyceride assay kit (Cat #E1103, Applygen Technologies).

### Confocal microscopy and statistical analysis

Images of adult fly brains were acquired by scanning a serial Z-stack of average 5-20 sections, each of 0.5, 0.8 or 0.125 μm thickness, using Zeiss LSM 700 (Oberkochen, Germany), Nikon A1 and C2 confocal microscope (Tokyo, Japan) with the 20X or 63X oil objective, respectively. The whole brain was positioned so that they can be scanned anteriorly to posteriorly (top to bottom). For statistical analyses of the mitochondrial area, original and unmodified images were imported into ImageJ (National Institutes of Health), and analyzed automatically using the Tubeness plugin. Mitochondrial volume was analyzed by Imaris (the Surface module) for 3D quantifications. For statistical analyses of LD number and intensity, original and unmodified images were imported into ImageJ, and the intensity threshold for the relevant channel was set so that maximum number of dots were selected without miscounting two adjacent dots into one. For colocalization measurements, primary images were imported into ImageJ, and analyzed automatically using the colocalization plugin with the intensity correlation tool, showing as Pearson’s correlation coefficient (R) or colocalized trace. All data were imported into GraphPad Prism 9 and shown in scatter plots or columnal bar graphs. For calculating the statistical significance, two-tailed unpaired t-test or ordinary one-way ANOVA followed by Tukey’s multiple comparisons test was used. P value less than 0.05 is considered significant. ns: no significance, p≥ 0.05; *: p<0.05; **: p<0.01; ***: p<0.001; ****: p<0.0001.

---

**Figure S1. *vamp7*-RNAi expression reduces *vamp7* mRNA levels and causes mitochondrial elongation in adult fly glia** (**A** and **B**) RT-qPCR analysis indicates that glial expression of *vamp7*-RNAi reduces the *vamp7* transcription levels significantly in both adult fly brains (RNAi^38300^ and RNAi^43543^, A) and FACS-sorted glia (RNAi^43543^, B). (**C** and **D**) Representative images (C,C’) and quantifications (D) of mitochondrial morphology in adult fly glia. Note that the average mitochondrial volume quantified by Imaris in 3D is significantly larger than the control glia when expressing *vamp7*-RNAi^43543^. All samples examined are 3-day-old adult fly brains unless noted otherwise. For RT-qPCR statistics, 20 independent fly brains for each genotype were analyzed. For FACS, GFP-positive glia cells from 200 independent fly brains for each genotype were collected. For single transgene experiment, *luc*-RNAi was used as a control. Scale bars of different sizes are indicated on the images. Serial confocal Z-stack sections were taken at similar planes across all genotypes, showing representative single-layer images. Statistical graphs are shown with scatter dots and the number of brains listed on top. Data are shown as mean ± SEM. P-values of significance (indicated with asterisks, ns no significance, * p<0.05, ** p<0.01, *** p<0.001 and **** p<0.0001) are calculated by ordinary one-way ANOVA followed by Tukey’s multiple comparisons test.

**Figure S2. VAMP7 regulates mitochondrial dynamics in glia independently of autophagosome-lysosome fusion and mitophagy** (**A** and **B**) Autophagic flux monitored by the dual reporter *UAS-GFP-mCherry-Atg8a* is impaired upon glial VAMP7 reduction. Note that the Pearson’s coefficient value analyzed by Imaris reveals the increased colocalization of GFP- and mCherry-positive signals when silencing *vamp7* expression in glia. (**C** and **D**) Representative quantifications (C) and images (D and D’) of mitochondrial morphology in glia with or without Bafilomycin A1 (BafA1). Note that no change in the mitochondrial area is observed when autophagosome-lysosome fusion is blocked by BafA1. (**E** and **F**) Glial STX17 inhibition fails to induce mitochondrial elongation. Note that the mitochondrial area remains unaffected when expressing *stx17*-RNAi in glia. (**G** and **H**) Representative images (H) and quantifications (G) of the DA neurons in 10-day-old flies expressing *luc-*RNAi, *vamp7-*RNAi^43543^, *stx17*-RNAi, or *snap29*-RNAi. Note that only vamp7-RNAi^43543^ expression causes DA neuron loss. (**I** and **J**) Representative quantifications (I) and images (J,J’) of mitochondrial morphology in glia when downregulating the expression of autophagy-related genes. Note that glial mitochondrial area remains largely intact when expressing *atg5*-RNAi or *ref(2)p*-RNAi, and smaller when expressing *atg1*-RNAi and *atg8*-RNAi. (**K-O**) Representative images (K) and quantifications (L-O) on mitochondria-lysosome contacts in fly glia expressing two reporters: *UAS-mCherry.mitoOMM* (labels the mitochondrial outer membrane, red) and *UAS-Lamp1.GFP* (labels lysosome, green). Contacts were co-labeled with the anti-Atg8a antibodies (Atg8a, magenta). Note a significant increase in the colocalization of mCherry- and GFP-positive signals upon glial VAMP7 reduction. Atg8a is minimally present in these dual-labeled contact sites, suggesting the absence of mitophagy. The selected areas in white squares in D,E,I are enlarged and aligned in D’,E’,I’. DA neurons are detected with anti-TH antibodies. All samples examined are brains of 3- or 10-day-old adult flies unless noted otherwise. For single transgene experiment, *luc*-RNAi was used as a control. Scale bars of different sizes are indicated on the images. A serial confocal Z-stack sections were taken at the similar plane across all genotypes, with representative merged images shown in A and H and single layer images shown in other panels. Statistical graphs are shown with scatter dots and the number of brains listed on top. Parameters for the colocalization and the mitochondrial area are quantified using Imaris and ImageJ as detailed in the Materials and Methods. Data are shown as mean ± SEM. P-values of significance (indicated with asterisks, ns no significance, * p<0.05, ** p<0.01, *** p<0.001 and **** p<0.0001) are calculated by two-tailed unpaired t-test between two sample groups and by one-way ANOVA compared among three or more sample groups followed by Tukey’s multiple comparisons test.

**Figure S3. Astrocytic VAMP7 reduction causes DA neuron loss** (**A-H**) Representative images (A, C, E, G) and quantifications (B, D, F, H) of the DA neurons in 10-day-old flies expressing *luc-*RNAi or *vamp7-*RNAi^43543^ using the different glial subtype drivers. Note that *vamp7*-RNAi^43543^ expression by the astrocyte driver GMR86E01-GAL4 causes decreased DA neuron number in the PPM1/2 cluster. DA neurons are detected with anti-TH antibodies. All samples examined are brains of 10-day-old adult flies unless noted otherwise. For single transgene experiment, *luc*-RNAi was used as a control. Scale bars of different sizes are indicated on the images. Serial confocal Z-stack sections were taken at similar planes across all genotypes, showing representative merged images. Statistical graphs are shown with scatter dots and the number of brains listed on top. Data are shown as mean ± SEM. P-values of significance (indicated with asterisks, ns no significance, * p<0.05, ** p<0.01, *** p<0.001 and **** p<0.0001) are calculated by ordinary one-way ANOVA followed by Tukey’s multiple comparisons test.

**Movie S1. *In-vivo* time lapse analysis of mitochondria-lysosome contact in the control flies.** Two reporters *UAS-mCherry.mitoOMM* (labels the mitochondrial outer membrane, red) and *UAS-Lamp1.GFP* (labels lysosome, green) are co-expressed to visualize the mitochondria-lysosome contacts. Note that mitochondria and lysosomes contact and dissociate over the imaging period. 5 images Z-stack sections were taken with 0.125 μm each at an interval of 10 seconds for 5 minutes. Merged images are shown. Video corresponds to Figure 2A. Scale bar, 1 μm.

**Movie S2-4. *In-vivo* time lapse analysis of mitochondria-lysosome contact in flies lacking glial VAMP7.** Two reporters *UAS-mCherry.mitoOMM* (labels the mitochondrial outer membrane, red) and *UAS-Lamp1.GFP* (labels lysosome, green) are co-expressed to visualize mitochondria-lysosome contact. A representative contact is shown (Movie S2, *repo>Lamp1GFP,mCherry.mitoOMM; Luc-*RNAi). Note the significant increase in colocalization of mitoOMM and Lamp1 upon glial VAMP7 reduction (Movie S3, *repo>Lamp1GFP,mCherry.mitoOMM; vamp7-*RNAi^38300^ and Movie S4, *repo>Lamp1GFP,mCherry.mitoOMM; vamp7-*RNAi^43543^). 5 images Z-stack sections were taken with 0.125 μm each at an interval of 10 seconds for 5 minutes. Merged images are shown. Videos correspond to Figure 2B. Scale bar, 1 μm.

**Movie S5-7. Glial VAMP7 regulates mitochondria lysosome contacts.** Two reporters *UAS-mitoOMM.mCherry* (labels the mitochondrial outer membrane, red) and *UAS-Lamp1.GFP* (labels lysosome, green) are co-expressed to visualize mitochondria-lysosome contact *in vivo*. A representative contact is shown (Movie S5, *repo>Lamp1GFP, mCherry.mitoOMM, LacZ; Luc-*RNAi). Note the significant increase in colocalization of mitoOMM and Lamp1 upon glial VAMP7 depletion (Movie S6, *repo>Lamp1GFP, mCherry.mitoOMM, LacZ; vamp7-*RNAi^43543^). Re-introduced 6xMyc-VAMP7 after decreased glial VAMP7 expression, mitochondria-lysosome contacts were rescued (Movie S7, *repo>Lamp1GFP, mCherry.mitoOMM, 6xMyc-vamp7; vamp7-*RNAi^43543^). 10 images Z-stack sections were taken with 0.125 μm each at an interval of 28 seconds for 5 minutes. Merged images are shown. Videos correspond to Figure 2E. Scale bar, 1 μm.

**Movie S8-12. Glial VAMP7 regulates mitochondria-lysosome contacts via Rab7 hydrolysis** Two reporters *UAS-mCherry.mitoOMM* (labels the mitochondrial outer membrane, red) and *UAS-Lamp1.GFP* (labels lysosome, green) are co-expressed to visualize mitochondria-lysosome contact. A representative contact is shown (Movie S8, *repo>Lamp1GFP, mCherry.mitoOMM, LacZ; Luc-*RNAi). Note the significant increase in colocalization of mitoOMM and Lamp1 upon glial VAMP7 depletion (Movie S9, *repo>Lamp1GFP, mCherry.mitoOMM, LacZ; vamp7-*RNAi^43543^). Co-expressing wild type Rab7 (*3xflag-rab7^wt^*) or GTP-bound Rab7 (*3xflag-rab7^Q67L^*) and *vamp7-*RNAi^435435^ formed a stable mitochondria-lysosome contact (Movie S10, *repo>Lamp1GFP, mCherry.mitoOMM, 3xflag-rab7^wt^; vamp7-*RNAi^43543^ and Movie S11, *repo>Lamp1GFP, mCherry.mitoOMM, 3xflag-rab7^Q67L^; vamp7-*RNAi^43543^). On the contrary, co-expressing GDP-bound Rab7 (*3xflag-rab7^T22N^*) and *vamp7-*RNAi^435435^ recovered the mitochondria-lysosome untethering (Movie S12, *repo>Lamp1GFP, mCherry.mitoOMM, 3xflag-rab7^T22N^; vamp7-*RNAi^43543^). 5 images Z-stack sections were taken with 0.125 μm each at an interval of 10 seconds for 5 minutes. Merged images are shown. Videos correspond to Figure 4A. Scale bar, 1 μm.

**Movie S13-16. Glial expression of TBC1D15-17 lacking the GAP activity disrupts mitochondria-lysosome contacts.** Two reporters *UAS-mCherry.mitoOMM* (labels the mitochondrial outer membrane, red) and *UAS-Lamp1.GFP* (labels lysosome, green) are co-expressed to visualize mitochondria-lysosome contact. A representative contact is shown (Movie S13, *repo>Lamp1GFP, mCherry.mitoOMM, 3xflag*). Note the significant increase in colocalization of mitoOMM and Lamp1 upon glial TBC1D15-17 lacking the GAP domain (Movie S16, *repo>Lamp1GFP, mCherry.mitoOMM, 3xFlag-TBC1D15-17*^Δ*GAP*^). Whereas expressing wild type Rab7 GAP TBC1D15-17^WT^ (Movie S14, *repo>Lamp1GFP, mCherry.mitoOMM, 3xFlag-TBC1D15-17^WT^*) or TBC1D15-17 lacking the disordered domain (Movie S15, *repo>Lamp1GFP, mCherry.mitoOMM, 3xFlag-TBC1D15-17*^Δ*disordered*^) in glia causes mitochondria and lysosome untethering successfully after contacts. 5 images Z-stack sections were taken with 0.395 μm each at an interval of 10 seconds for 5 minutes. Merged images are shown. Videos correspond to Figure 5I. Scale bar, 1 μm.

**Movie S17-21. VAMP7 regulates mitochondria-lysosome contacts in glia via the Rab7 GAP TBC1D15-17.** Two reporters *UAS-mCherry.mitoOMM* (labels the mitochondrial outer membrane, red) and *UAS-Lamp1.GFP* (labels lysosome, green) are co-expressed to visualize mitochondria-lysosome contact. A representative contact is shown (Movie S17, *repo>Lamp1GFP, mCherry.mitoOMM, LacZ; Luc-*RNAi). Note the significant increase in colocalization of mitoOMM and Lamp1 upon glial VAMP7 depletion (Movie S18, *repo>Lamp1GFP, mCherry.mitoOMM, LacZ; vamp7-*RNAi^43543^). Co-expressing *vamp7-*RNAi^43543^ and wild type Rab7 GAP 3xflag-TBC1D15-17^WT^ (Movie S19, *repo>Lamp1GFP, mCherry.mitoOMM, 3xFlag-TBC1D15-17^WT^; vamp7-*RNAi^43543^) or TBC1D15-17 lacking the disordered domain (Movie S20, *repo>Lamp1GFP, mCherry.mitoOMM, 3xFlag-TBC1D15-17*^Δ*disordered*^*; vamp7-*RNAi^43543^) causes mitochondria and lysosome untethering successfully after contacts. Whereas co-expressing glial *vamp7*-RNAi and TBC1D15-17 lacking the GAP domain (Movie S16, *repo>Lamp1GFP, mCherry.mitoOMM, 3xFlag-TBC1D15-17*^Δ*GAP*^*; vamp7-*RNAi) inhibits untethering after mito.OMM and Lamp1.GFP contacts. 5 images Z-stack sections were taken with 0.395 μm each at an interval of 10 seconds for 5 minutes. Merged images are shown. Videos correspond to Figure 6A. Scale bar, 1 μm.

