## Supplementary figures and images for "VAMP7-dependent mitochondria-lysosome contacts contribute to glial mitochondrial dynamics and dopaminergic neuron survival"

### Figure S1

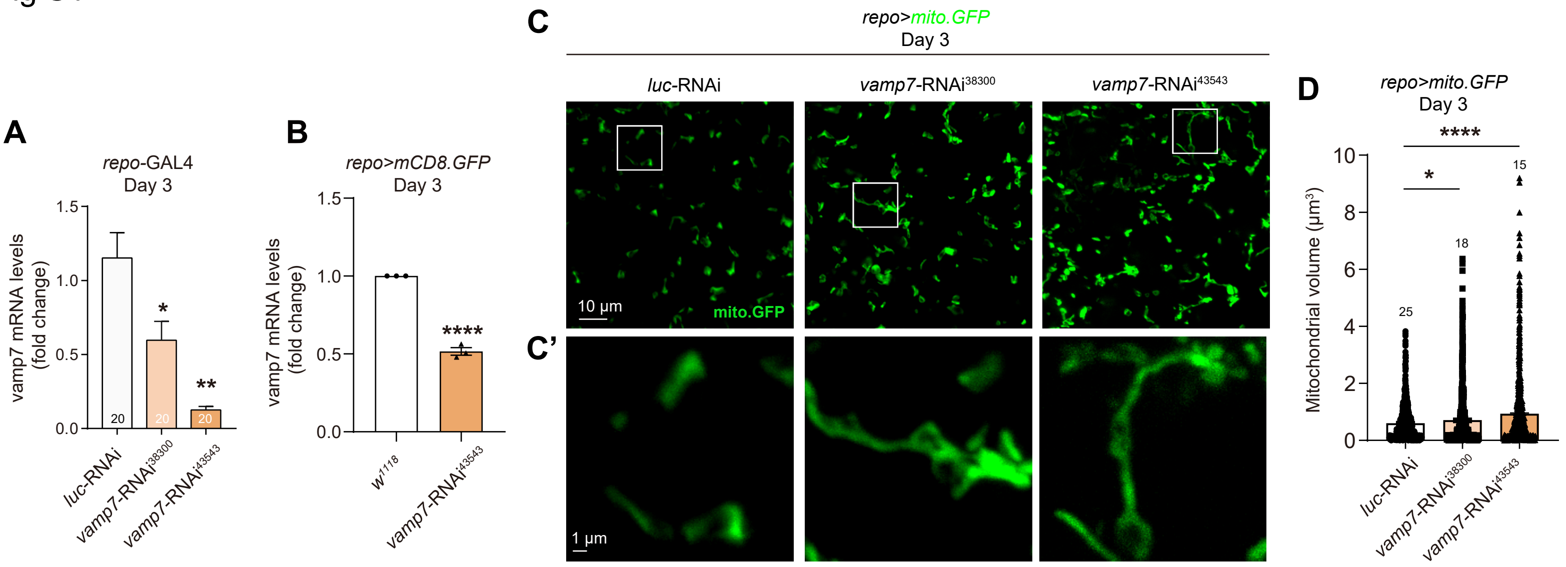

### Figure S2

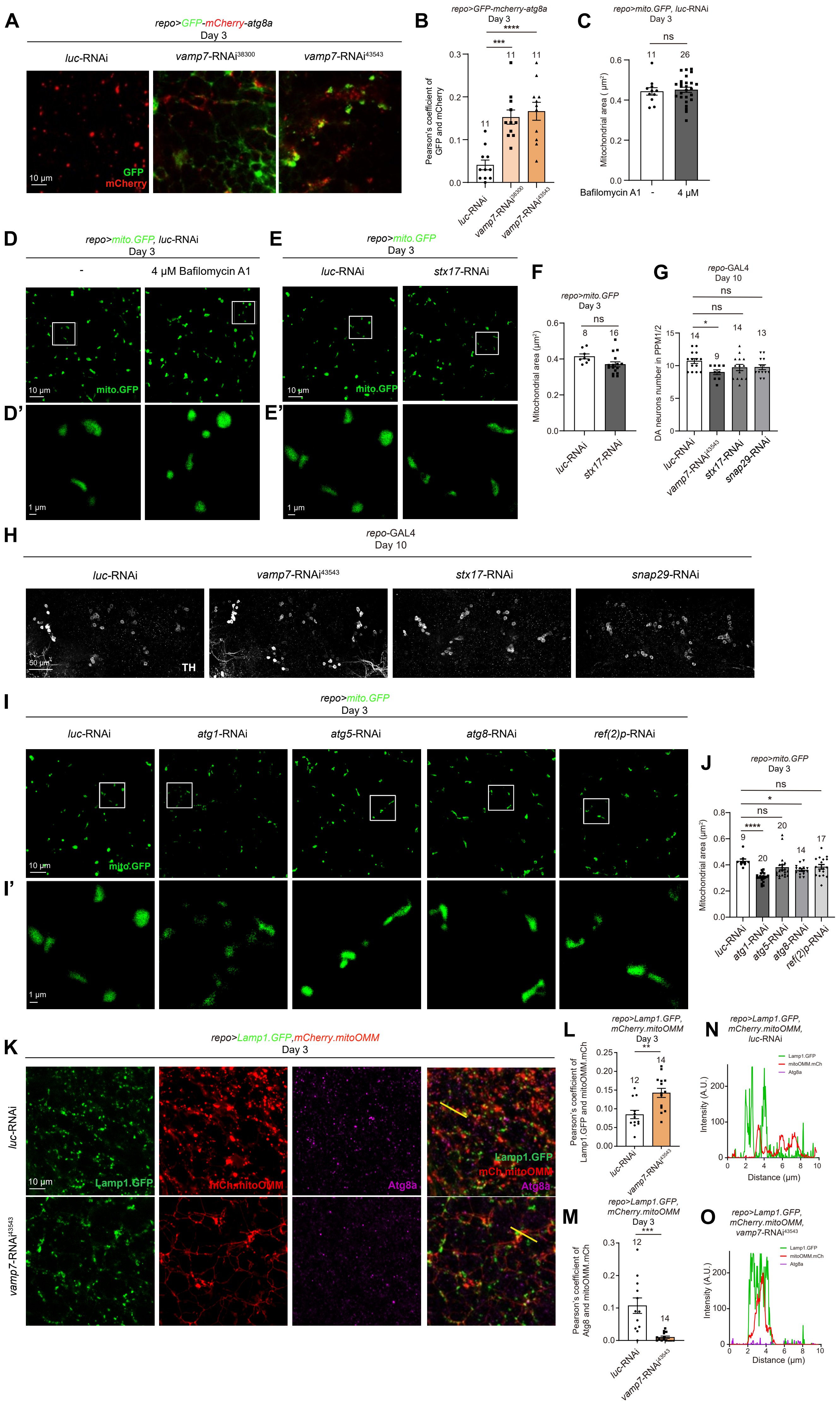

### Figure S3

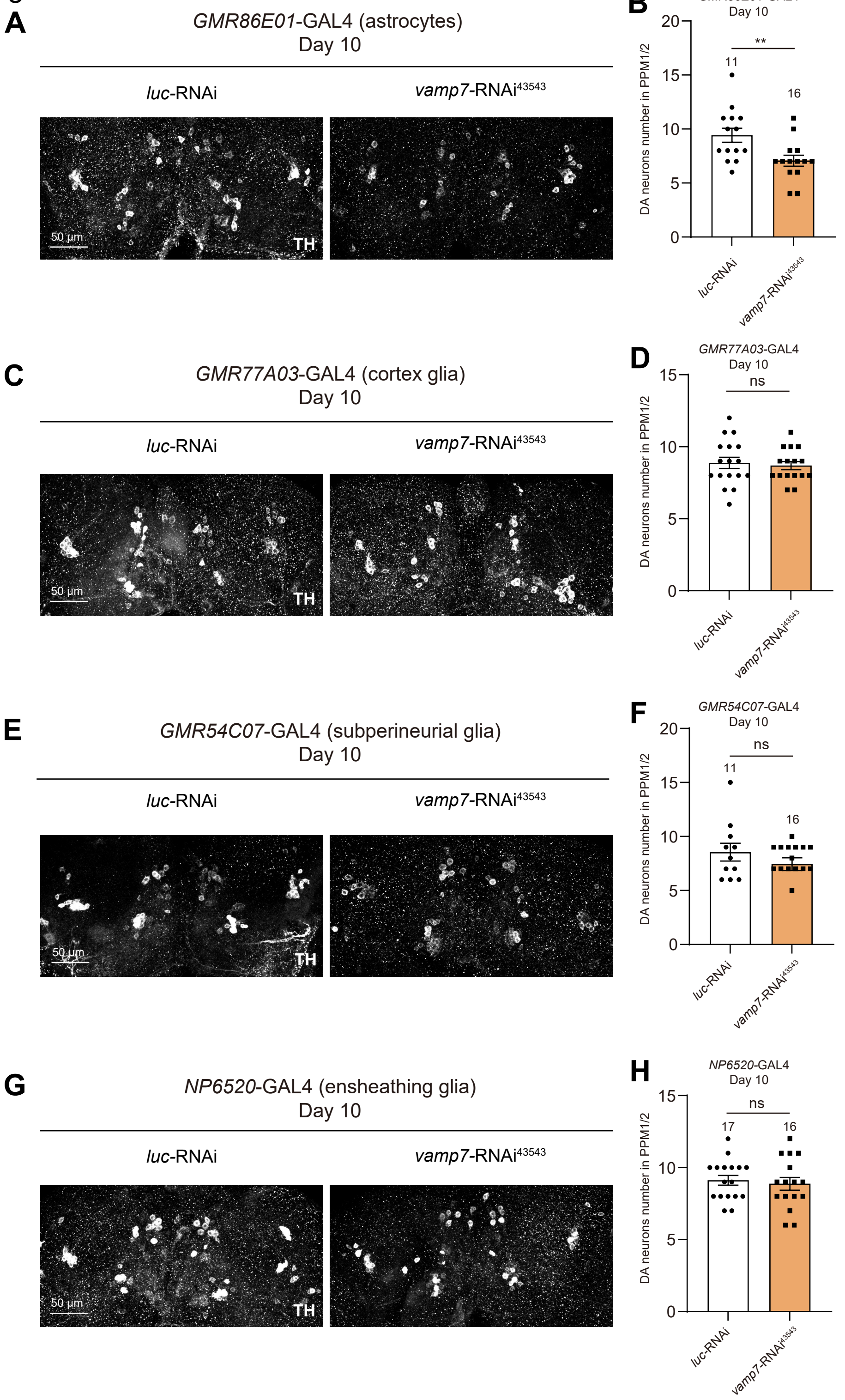
